# Investigating endometriosis development and endometriosis-related pain over time and in relation to estrogen in a laparoscopic mouse model

**DOI:** 10.1101/2024.03.28.583957

**Authors:** Daniëlle Peterse, Alejandra Verhassel, Amelie Fassbender, F. O Dorien, Arne Vanhie, Anne-Sophie Van Rompuy, Philippa Saunders, Joris Vriens, Thomas M. D’Hooghe

**Author notes:** Correspondence: Corresponding author: Thomas M. D’Hooghe, M.D., Ph.D.; Faculty of Medicine, Leuven University, Belgium; Department of Development and Regeneration, Herestraat 49, 3000 Leuven, Belgium. Harvard Medical School, Department of Surgery; Boston Children’s Hospital, Vascular Biology Program, Boston, MA 02115, USA.

## Abstract

**Background:** Endometriosis is a complex disease, and its pathophysiology is still unclear. Therefore, endometriosis animal models need to be carefully selected and examined to be useful for identification of novel therapies for women with endometriosis. In this study, we evaluated endometriosis-associated pain, and time- and estrogen-related development of endometriotic lesions after laparoscopic implantation of menstrual endometrium in a homologous mouse model for endometriosis.

**Methods:** Endometriosis was induced by laparoscopic introduction of 10 menstrual endometrial tissue pieces into the peritoneum of ovariectomized recipient mice (59 estrogen-substituted; 59 estrogen-depleted). Sham animals (57 estrogen-substituted; 60 estrogen-depleted) received 10 pieces of perigonadal adipose tissue. The animals were sacrificed at 1, 2, 3, 4, 6 or 8 weeks after induction, the attached peritoneal implants localized and excised and immunohistochemically analyzed. Additionally, endometriosis-related pain was evaluated by measuring mechanical allodynia, thermal hyperalgesia, locomotor activity and anxiety-like behavior before and after tissue implantation.

**Results:** At least one implant per mouse could be retrieved in 94% (111/118) of the endometrial tissue animals and in 78% (91/117) of the adipose tissue animals (p<0.001). Peritoneal implant take rate was significantly higher in endometrial tissue animals (2.5±1.4) compared to adipose tissue animals (1.6±1.5) (p<0.0001), regardless of estrogen supplementation and time of sacrifice. Hemosiderin could be observed more often (p<0.0001) in attached peritoneal implants of the endometrial tissue animals (67%, 68/101), compared to the adipose tissue animals (37%, 31/83). Ki67 staining showed a higher proliferation index in the attached peritoneal implants retrieved after one week, compared to the other time points of both endometrial tissue and adipose tissue animals. The behavioral test showed no significant difference in mechanical and thermal sensitivity, locomotor activity and anxiety-behavior between the menstrual endometrial tissue and adipose tissue implanted animals. Nevertheless, the estrogen-substituted animals showed decreased activity in the tests featuring thermal nociception and anxiety-like behavior, compared to the estrogen-depleted animals. Additionally, time after implantation showed to have a positive effect on thermal sensitivity, locomotor activity and anxiety-related behavior in all animals, as the mice became less sensitive to thermal stimuli, more active in the open field test and buried less marbles in the marble burying test.

**Conclusion:** This study showed an increased attachment of menstrual endometrium compared to adipose tissue in the peritoneum when using laparoscopic induction. There was no apparent influence of estrogen on tissue attachment, proliferation or appearance. A decrease in cell proliferation in peritoneal implants occurred over time. Locomotor activity, anxiety-like behavior, and mechanical and thermal sensitivity of the animals was not affected after induction of endometriosis, regardless of the type of implanted tissue. Altogether, we showed that the current methodology used to induce endometriosis was not sufficient to develop endometriotic lesions that contained both stromal and epithelial cells. Moreover, the current methodology was not able to detect specific endometriosis-related pain.

**GRAPHICAL ABSTRACT:** 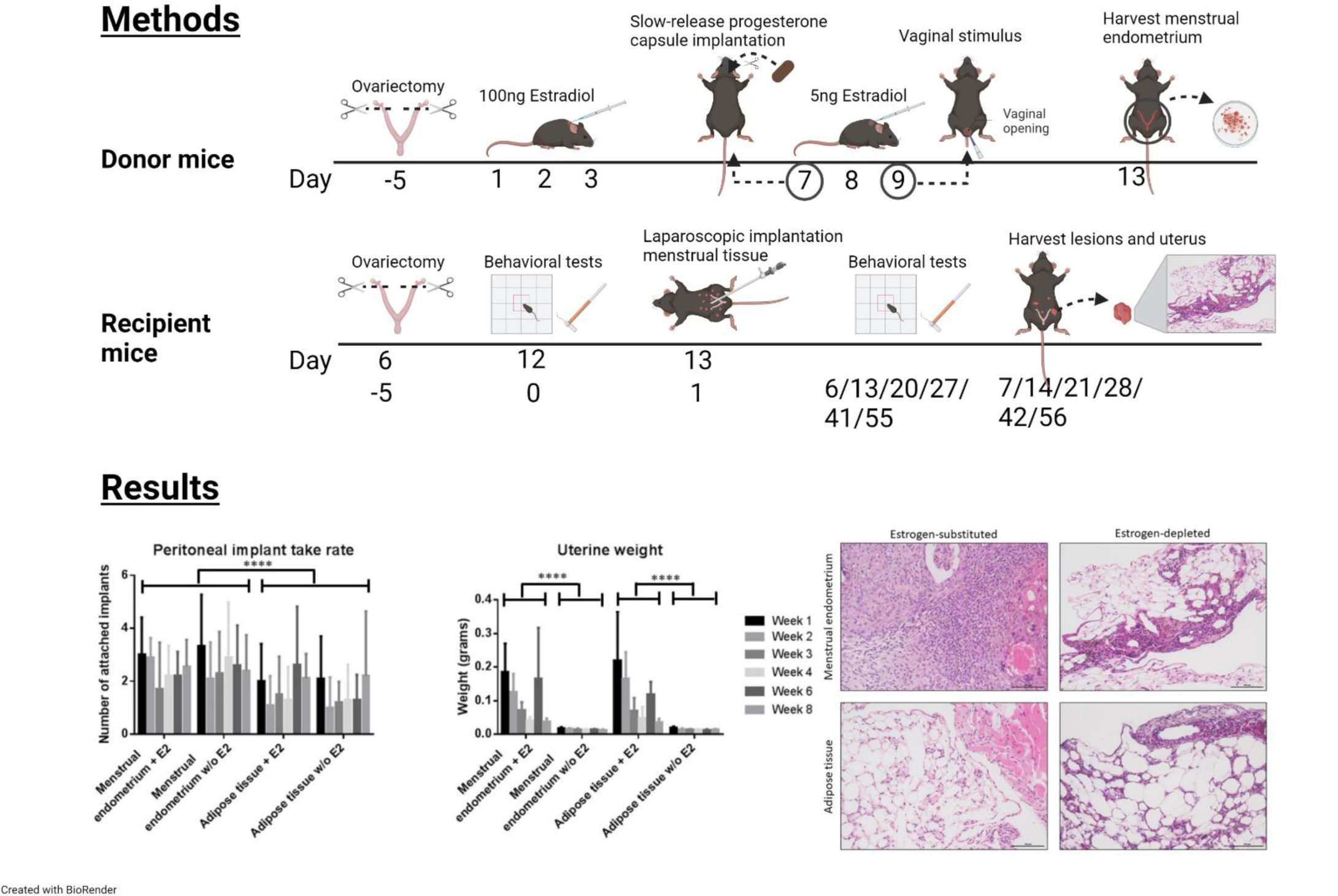

## 1. INTRODUCTION

Endometriosis is an estrogen-dependent gynecological disease, characterized by endometrial-like tissue outside the uterus and is associated with pain and infertility^1^. The origin of endometriosis is still not clear, and several theories to explain its pathophysiology have been proposed. The most accepted theory is Sampson’s theory of retrograde menstruation, attributing the development of endometriosis by pelvic attachment of shed endometrial tissue, which flows backwards through the fallopian tubes during menstruation^1, 2^.

Estrogen plays a key role in the development of endometriosis, as high estrogen levels have been found near lesions and -in less abundant amounts-in the endometrium^3, 4^ and the severity of endometriosis-related symptoms can be reduced by suppressing ovarian function^5–7^.

Dysmenorrhea (pain during menstruation), dyspareunia (pain during sexual intercourse), dysuria (pain during urination) and chronic pelvic pain are the most common symptoms of endometriosis related pain^8^. Different mechanisms have been described to explain the origin of endometriosis related pelvic pain. Firstly, the unmyelinated sensory C fibers, myelinated sensory Aδ fibers and adrenergic nerve fibers, which are present in and around endometriotic lesions^9, 10^, function as nociceptors. Through the presence of these nerve fibers, endometriosis can act as noxious stimulus and trigger transduction, most likely via neurotrophin-induced signalling, although the exact mechanism remains unclear^9, 10^. This mechanism is supported by a correlation between nerve innervation in peritoneal lesions and endometriosis-related pain^11^. Secondly, endometriosis related pain can have an inflammatory origin, due to the chronic inflammation caused by endometriosis. Several inflammatory factors, such as IL-1, IL-6, IL-8, and nerve growth factor have shown to be increased in the peritoneal fluid or ectopic lesions of endometriosis patients. Lastly, endometriosis associated pain can be neuropathic, caused by endometriotic lesions affecting the sensory component of the nervous system^9, 12^. Consequences of neuropathic pain are sensitization of the central and peripheral nervous system. It has been described that endometriosis patients experience chronic pelvic pain, caused by central sensitization^13^. Furthermore, these patients show increased allodynia in all body parts, compared to healthy controls^9^. This can be explained by the increased expression of the nociceptors TRPV1 and TRPA1 in the peritoneum and endometriotic lesions of women with endometriosis^14–16^.

Spontaneous endometriosis has only been described in humans and non-human primates^17^. For this reason, it is believed that the baboon model for endometriosis closely mimics human disease regarding etiology and symptoms ^18^. Nevertheless, the use of non-human primates is restricted due to limited availability, high costs and ethical reasons^19^.

Rodent models are generally attractive models to use for pre-clinical studies due to their low cost and ease of manipulation^20, 21^. However, most laboratory rodents have a four-day estrous cycle without spontaneous decidualization or menstruation, and therefore do not develop endometriosis spontaneously. To overcome this limitation, the menstruating mouse model, as first described by Brasted *et al.*^22^ and refined by Greaves *et al.*^23^, has been adapted for endometriosis research and shown to result in lesions that mimic the phenotype of those in patients^23, 24^. Nevertheless, the time-related development and progression of the endometriotic lesions, including neurogenesis and lymphangiogenesis, after implantation of murine menstrual endometrium has never been studied before, although it is important that mouse models are well characterized and validated before they can be used for the identification of new potential therapies for endometriosis treatment^25, 26^.

In mice, estrogen has been shown to be involved in lesion attachment in natural cycling C57Bl/6J mice, but not in BALB/c mice, although no estrogen-dependent differences in lesion size, type or location could be identified^27, 28^. In ovariectomized and estrogen-substituted endometriosis models, estrogen is not required for lesion establishment, although ER-α plays a leading role in the attachment of endometrial tissue^29^. Moreover, it has been shown in mice that estrogen signaling via ER-α mediates lesion growth by increasing lesion size and proliferation of epithelial cells^29^. Furthermore, there is evidence that estrogen plays a role in the maintenance of the implanted lesions, although contradictory results have been published^29–36^.

Clinically, it is not ethical to perform repeated laparoscopies in endometriosis patients to study progression of the disease. Therefore, performing longitudinal studies to investigate both lesion progression and regression would highly increase the translational value of pre-clinical endometriosis models^26^. Nevertheless, only a limited number of studies performed in endometriosis mouse models evaluate both induction and regression of the disease, although a gradual resolution of lesions has been identified^26, 38^. Recently, it has been shown that only 23% of studies that used mouse models to investigate endometriosis included timepoints beyond four weeks after induction, although it has been demonstrated that endometriosis lesions can gradually resolve over time^37, 38^. This indicates that more longitudinal studies are needed to investigate the natural life cycle of lesions and the effect of new therapies on lesion attachment, development and regression^26^.

A crucial step in development and validation of the mouse model for endometriosis induced with menstrual endometrium requires the demonstration of endometriosis-associated pain and hypersensitivity^8^. So far, only 2-3% of publications have evaluated a spontaneous (non-evoked) pain in mouse models for endometriosis, although the many researchers believe that measuring spontaneous pain is more comparable to the pain that women with endometriosis experience^39, 40^.

Additionally, neurogenesis should be studied in endometriosis animal models to gain knowledge regarding the poorly-understood mechanisms behind endometriosis-associated pain^25^. It has been shown that nerve fibers are present in the lesions of different mouse models for endometriosis^41, 42^, and that nerve fiber density does not decrease after estrogen depletion for one week^43^. Nevertheless, information regarding nerve growth in relation to presence or absence of estrogen over a longer time in endometriosis mouse models is lacking.

Therefore, we wanted to investigate lesion ontogenesis, endometriosis-related neurogenesis and endometriosis associated pain symptoms in a novel mouse model for endometriosis in relation to normal estrogenic and hypo-estrogenic conditions at several time points.

## 2. MATERIALS AND METHODS

### 2.1. Animals

A total of 353 eight-week-old female C57BL/6JRj were ordered from Janvier (Saint Berthevin Cedex, France). Five animals were housed per cage, with a room temperature of 23±1.5°C, humidity between 40-60%, and a 14:10 hour light-dark cycle. The mice were fed *ad libitum* with food pellets and water. All animal studies are approved by the ethical committee (ethical approval number: P031/2013).

### 2.2. Randomization

The animals were divided in 60 donor mice and 240 recipient mice, using computerized randomization. The donor mice were used to obtain menstrual endometrium and adipose tissue for intra-abdominal placement in recipient mice.

The 240 recipient mice were subdivided into four groups (each consisting of 60 mice) depending on the type of implanted tissue (menstrual endometrium or adipose tissue) and estrogen-substitution or -depletion. Each of these four groups were divided into six subgroups (10 mice per subgroup), depending on the timing of sacrifice, which was 1, 2, 3, 4, 6, or 8 weeks after induction of endometrial- or adipose tissue, see Figure 1 for a detailed timeline regarding the proceedings of the donor- and recipient animals. The number of recipient mice per group (n = 10) was based on a similar number used in a study evaluating pain in a mouse model for endometriosis^44^.

**Figure 1:**
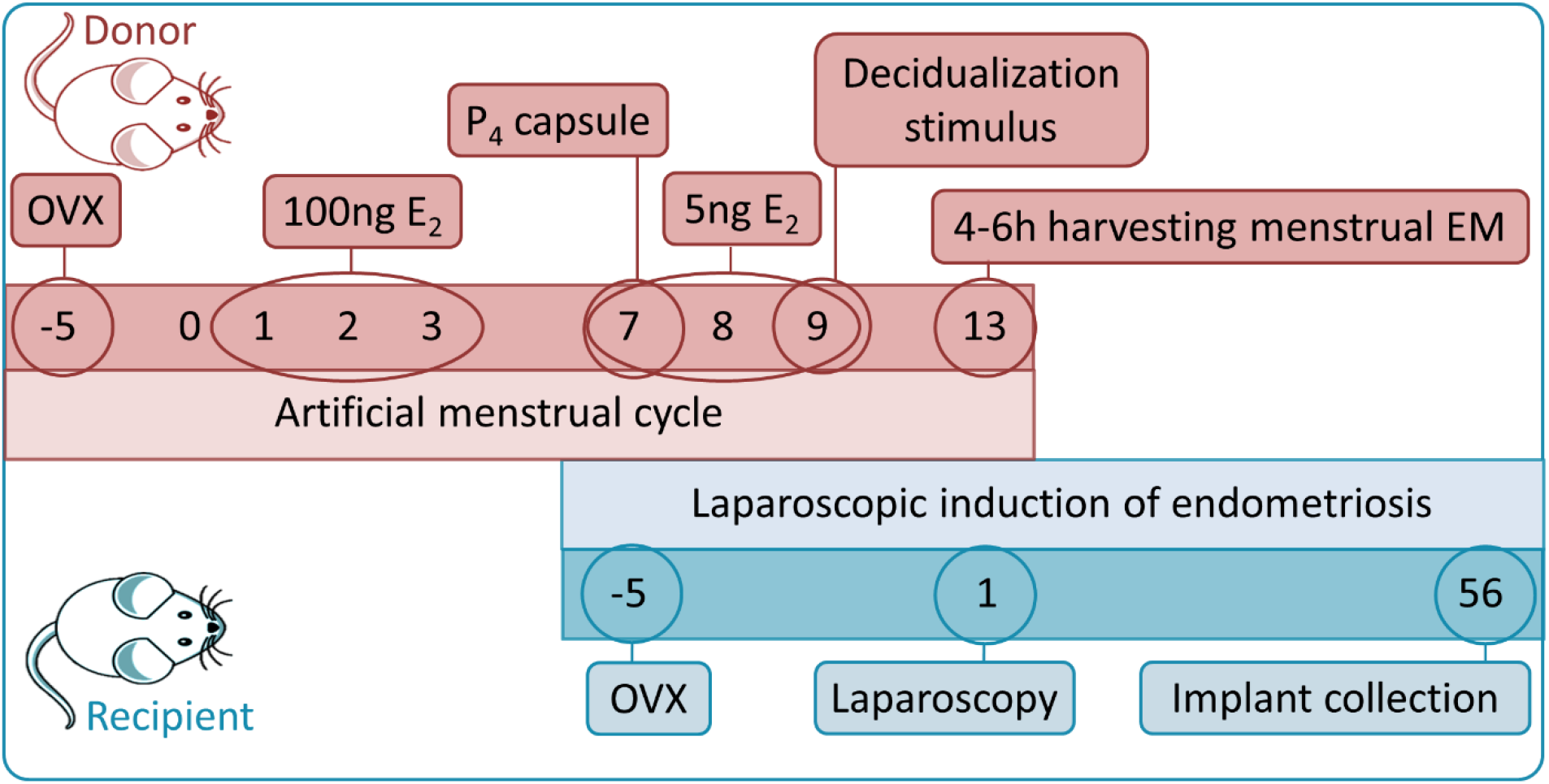
Schematic presentation of the treatment schedules for donor and recipient mice.

Finally, 306 animals were included in the study; 71 donor mice and 235 recipient animals. Moreover, 47 animals were excluded from all analyses for the following reasons: death due to complications during ovariectomy (n = 3), complications during laparoscopy (n = 12), unknown reasons (n = 3), presence of infections (n = 6), or peritoneal malformations observed at sacrifice (n = 5). A total of 18 recipient mice were excluded due to difficulties with the implanted estrogen capsule, as the capsule did not release a sufficient amount of estrogen, based on the results of the vaginal smears.

### 2.3. Anesthesia

Isoflurane-induced anesthesia (3% Iso-Vet; Eurovet, Bladel, The Netherlands) was used in both donor- and recipient mice to perform ovariectomy, implantation, renewal and removal of the slow-release progesterone- and estrogen capsules, decidualization induction, and exsanguination. Prior to the surgical procedure, the appropriate area was shaved and disinfected using an iodine-based solution (BDH Prolabo, VWR International, Heverlee, Belgium). After surgery, the skin incision was sutured with a 6-0 polyglycolic acid thread (Dexon II, Davis & Geck, Gosport, UK) and a subcutaneous injection of 0.05 mg/kg buprenorphine (Vetergesic, Ecuphar, Breda, The Netherlands) diluted in saline (0.9% NaCl; Viaflo, Baxter, Utrecht, The Netherlands) was given on two consecutive days as painkiller.

For laparoscopy, animals were anesthetized with an intraperitoneal injection using a mixture of 80 mg/kg ketamine (Nimatek, Eurovet Animal Health, Bladel, The Netherlands), and 10 mg/kg xylazine (XYL-M, VMD nv, Arendonk, Belgium).

### 2.4. Ovariectomy

To exclude aberrations in the hormonal environment due to the endogenous hormone production, all donor and recipient animals were ovariectomized. Briefly, a dorsal midline skin incision was made caudal to the posterior border of the ribs. Subsequently, the peritoneum was incised at the level of the ovaries, and the ovaries were removed. Hereafter, the skin was sutured, and the animals allowed to recover for one week.

### 2.5. Donor mice

#### 2.5.1. Retrieval of menstruating endometrium and adipose tissue

Donor mice have been treated as described previously^24, 45^. In short, ovariectomized donor mice were subcutaneously injected with 100 ng estradiol for three consecutive days. After a three-day resting period, 5 ng estradiol was injected subcutaneously for three consecutive days. On the first day of this second series of estrogen injections, a progesterone pellet (5 mg) was implanted, and an intra-uterine decidualization stimulus, using 100 µl of sesame oil with an additional mechanical stimulus was applied via the vagina in both uterine horns on the last day. Four days after the decidualization stimulus, the progesterone pellet was removed and 4-6 hours later, the animals were anesthetized, and the menstrual endometrium and adipose tissue were harvested, after which the animals were sacrificed by exsanguination.

#### 2.5.2. Tissue processing

The harvested uterus was longitudinally opened, the dissociating menstrual endometrium was gently scraped from the myometrium and weighed on a microgram balance. This menstrual tissue was punched into 1-2 mm diameter pieces by using a biopsy puncher, and 10 pieces were selected for each recipient animal. Furthermore, 1-2 mm diameter pieces of perigonadal adipose tissue were punched, using a biopsy puncher, and 10 pieces were selected for each recipient animal. We choose for perigonadal adipose tissue in this study for our sham animals, since this has been used in different studies describing animal models for endometriosis and does not attach onto the peritoneal wall or visceral organs when injected.^46–48^

### 2.6. Recipient mice

#### 2.6.1. Vaginal smear

The estrous cycle of the animals was staged by vaginal cytology, daily before ovariectomy and once a week after this surgery. This was performed to check if the ovariectomy was successful, to ensure that the implanted estrogen capsule was working properly and to determine the estrous cycle phase during behavioral tests, as it has been proposed that differences in estrogen levels can have an effect on animal behavior. The estrogen levels of the animals were determined using vaginal cytology, since this was less invasive for the animals compared to repetitive blood sampling.

#### 2.6.2. Laparoscopic induction of recipient mice with menstrual endometrium or perigonadal adipose tissue

Laparoscopic induction of endometriosis was performed in all recipient mice. Inducing endometriosis via laparoscopy has the advantage that endometriosis lesions will develop at places similar to where lesions typically are found in endometriosis patients. In addition, it has been shown that laparoscopy results in a higher peritoneal implant take rate, compared to blind peritoneal injection^49^.

To perform laparoscopy, animals were anesthetized, secured in a supine position, and intubated endotracheally with a 20G catheter, which was connected to a mechanical respirator (Rodent Ventilator; Harvard Apparatus, Holliston, MA, USA) using 250 μl air/stroke and 180 stokes/minute.

Through a 3.5 mm abdominal incision, the laparoscope (Karl Storz, Tüttlingen, Germany was placed in the peritoneal cavity, as described before^24, 49^, with a pressure of 15 mmHg. Hereafter, 10 menstrual endometrial fragments (diameter 1-2 mm) from donor animals were inserted using two 14G catheters (Insyte, Vialon, BD, Madrid, Spain) under laparoscopic guidance in both left and right flank and behind the uterus. Sham induction was done by laparoscopic insertion of perigonadal adipose tissue fragments at the same locations.

### 2.7. Estrogen supplementation

Immediately after finishing the laparoscopy, a silastic capsule containing estrogen was implanted subcutaneously in the estrogen-substituted animals, in order to restore the circulating estrogen levels. A subcutanous slow-release capsule to supplement circulating estrogen in the animals was chosen over daily oral or subcutanous estrogen injections, since slow-release capsules avoid daily hormonal fluctuations as well as reduces stress related to daily animal handling. The capsules were prepared as described by Ingberg, *et al.*^50^. In short, 1 cm of Silastic tubing (Dow Corning, Seneffe, Belgium) was filled with 17β-estradiol (Sigma-Aldrich, Dielegem, Belgium) dissolved in ethanol and diluted in arachnid oil to a final concentration of 36 μg/ml. The estrogen capsule was replaced four weeks after tissue implantation in the estrogen-substituted animals that would be sacrificed six or eight weeks after laparoscopy. The slow-release estrogen capsule was replaced since it has been shown that the amount of released estrogen is gradually decreasing over time^50^.

### 2.8. Sample collection

The animals were sacrificed by exsanguination at pre-defined time points. Laparotomy was performed using a stereomicroscope (Zeiss, Oberkochen, Germany) to record and photograph attached peritoneal implants with respect to number, length, localization and color. Moreover, macroscopically visible attached peritoneal implants, blood, peritoneum, and uterus were collected. Due to the small size of the attached peritoneal implants, the tissue was excised together with the surrounding tissue. Removed attached peritoneal implants and uterus were weighed. Part of the peritoneum, uterus, and endometrial-like and adipose tissue lesions were fixed in 4% formaldehyde for paraffin embedding.

### 2.9. Quantification of attached peritoneal implants

The incidence of tissue attachment was defined as the percentage of animals in which at least one attached implant could be retrieved. Moreover, the implant take rate was defined as the number of retrieved lesions divided by the number of implanted fragments (10 per animal).

### 2.10. Histology and immunohistochemistry

From each animal with three or more lesions, the lesion with the highest weight was chosen for immunohistochemistry (IHC). Paraffin-embedded attached peritoneal implants were cut into 4 µm sections and the sections were stained with hematoxylin and eosin (H&E), to visualize the structure of the lesion. Thereafter, IHC was performed for Ki67; vimentin; cytokeratin; Estrogen Receptor-α; Progesterone Receptor; Podoplanin; and Pgp9.5. Relevant positive and negative controls were used for each staining, see Table 1 for information regarding the positive controls, while buffer controls were used as negative control.

**Table 1:** Type of stimulus and reaction for the different applied behavioral tests^61^.

| Test | Pain stimulus | Type of pain/behavior | Caused by |
| --- | --- | --- | --- |
| Hot water test | Evoked pain | Thermal hypersensitivity/hyperalgesia | Central sensitization (inflammatory or neuropathic pain) |
| Acetone test | Evoked pain | Thermal hypersensitivity/hyperalgesia | Central sensitization (inflammatory or neuropathic pain) |
| Hargreaves test | Evoked pain | Thermal hypersensitivity/hyperalgesia | Central sensitization (inflammatory or neuropathic pain) |
| Von Frey Hair test | Evoked pain | Mechanical allodynia | Central sensitization (inflammatory or neuropathic pain) |
| Open field test | Spontaneous pain | Spontaneous behavior | Nociceptive, inflammatory or neuropathic pain |
| Marble burying test | Affective and emotional consequences of pain | Anxiety-like behavior | Nociceptive, inflammatory or neuropathic pain |

In short, the samples were deparaffinized and endogenous peroxidase was blocked using 3% H2O2, followed by antigen retrieval for one hour at 90°C. Thereafter, blocking buffer was applied to reduce non-specific staining and the samples were incubated with primary antibody (see Supplementary Table 1 for details). Next, the blocking buffer was applied again, followed by a 30-minute incubation step with biotinylated secondary antibodies at room temperature. Finally, streptavidin/horseradish peroxidase was applied onto the samples for 30 minutes, after which the receptors were visualized.

Each stained cross-section of the attached peritoneal implants was analyzed using a Nikon Eclipse Ci Microscope (Nikon, Brussels, Belgium) equipped with a Nikon DS-Fi3 camera.

Gross morphology of the attached peritoneal implants and presence of hemosiderin were visualized using H&E staining. Attached peritoneal implants were considered endometriotic if they showed histological presence of both endometrial glands and stroma. Hemosiderin, which is an iron complex most commonly found in hemosiderin-laden macrophages, is considered to be a marker for endometriosis^51, 52^, due to a disrupted iron metabolism in endometriosis patients^53^.

Vimentin and cytokeratin were analyzed in a binary way (as either being present or absent in the retrieved peritoneal implants). For PGP9.5 and Podoplanin, the number of nerve fibers or lymph vessels per mm^2^ was calculated. ERα and PR were analyzed using the Allred scoring system, which is commonly used for breast cancer analyses^54^. The Allred scoring system grants a score for the number of positive stained cells, ranging from 0 (negative) to 5 (100% positive). Next, staining intensity was rated from 0 (negative) to 3 (intense staining). Both scores were added up and this resulted in a scoring system varying from 0-8.

For Ki67, the proliferation index was calculated by counting the number of positive cells per total of 100 cells. For each slide, this calculation was performed at three random places and the average number of positive cells was used for analysis.

### 2.11. Pain and behavioral tests

All pain and behavioral tests were performed before ovariectomy, preferably when the animals were in the estrus phase of the estrous cycle, since estrogen levels, both in natural cycling and estrogen-substituted animals, are known to influence animal behavior^55–60^. Animals were subjected to a second set of behavior experiments at the end of the study, one day before the animals were sacrificed. The stimulus and type of reaction of the different pain and behavioral tests are summarized in Table 1.

### 2.12. Thermal sensitivity

To test cold hypersensitivity, nocifensive behavior (response to pain or discomfort), such as total reaction time and number of times the animals exhibited abdominal grooming behavior, was measured for a total of 2 minutes after pipetting 50 µl acetone onto the shaved abdomen of the animals^62, 63^. To test heat hyperalgesia, the hot water test was performed. In this test, 50 µl polyethylene glycol 400 (PEG) of 50°C was applied onto the shaved lower abdomen of the animals. Nocifensive behavior was analyzed measuring latency to first response, total response time, and abdominal grooming for a total of 2 minutes.

Furthermore, the Hargreaves test was performed onto the shaved abdomen to measure sensitivity to an infrared heat stimulus, by measuring withdrawal time^64^.

### 2.13. Mechanical sensitivity

Nociception to mechanical stimuli was measured using the Von Frey hair test (simplified up-down method), which was applied onto the shaved midline area of the lower abdomen^65, 66^.

### 2.14. Psycho-physiological tests

#### 2.14.1. Marble burying test

In the marble burying test, 20 marbles were evenly spaced in a standard cage containing 125 gr-130 gr bedding material and each animal was exposed for 30 min to the marbles. At 5, 10, 15, 20, 25 and 30 min after exposure, the number of buried marbles (at least 2/3 covered by dust) was counted^67^.

#### 2.14.2. Voluntary locomotor activity (open field test)

In the open field test, mice could move around freely in an open field (50 x 50 cm) for 5 minutes, while spontaneous locomotor activity was measured (Bioseb, France).

### 2.15. Statistical analysis

Statistical analysis was performed using GraphPad Prism 6 software and SAS software (version 9.4 of the SAS System for Windows), and a p-value <0.05 was significant.

Uterine weight, number of lesions, and results from the immunohistochemistry were compared between the adipose tissue groups and the endometrial tissue groups and between the estrogen-supplemented and estrogen-depleted groups using two-way ANOVA, Chi-square or Fisher’s exact test.

Data analysis for pain behavior was performed using linear models with the change in pain score (after - before) as response variable, and treatment (endometrial tissue or adipose tissue), estrogen (substituted or depleted), and weeks (after implantation) as explanatory variables. Model construction was performed by including all two-way and three-way interactions, and hierarchically excluding non-significant interaction terms from the model. The final model was then the model with all main effects and significant interaction effects.

For the marble burying test, analysis was based on the longitudinal data. A random effect for animal was included to correct for clustering. The minute of measurement was additionally added to the fixed-effects model, and model construction was performed analogously as previously described.

## 3. RESULTS

### 3.1. Donor mice

A total of 71 donor mice were used to obtain menstrual endometrium and perigonadal adipose tissue for laparoscopic implantation into recipient mice. The average uterine weight of the decidualized uteri of the donor animals was 289±99 mg and ranged between 105 mg till 517 mg.

Overall, 30% (18/60) of the mice showed full bicornuate decidualization, while 35% (21/60) showed full decidualization in one horn and partial decidualization in the other uterine horn. Furthermore, in 15% (9/60) of the animals, both uterine horns were partially decidualized. Finally, 18.3% (11/60) of the mice showed unicornuate decidualization and in 1.7% (1/60) of the animals, no decidualization could be observed at all. Details regarding number of decidualized horns were not noted in 11 animals.

### 3.2. Recipient mice

#### 3.2.1. Tissue implantation

All animals that survived till the end of the study, 118 endometrial tissue mice and 117 adipose tissue animals, were included in the analyses. From the 118 endometrial tissue mice, 94% (111/118 animals) presented with at least one attached lesion, which was significantly higher (p<0.001) compared to adipose tissue animals, of which 78% (91/117 animals) showed at least one attached implant (Table 2).

**Table 2:** Percentage of recipient animals showing at least one attached peritoneal implant (incidence)

|  | Endometrial tissue + E<sub>2</sub> | Endometrial tissue – E<sub>2</sub> | Adipose tissue + E<sub>2</sub> | Adipose tissue – E<sub>2</sub> |
| --- | --- | --- | --- | --- |
| Week 1 | 100% (10/10) | 100% (9/9) | 90% (9/10) | 100% (8/10) |
| Week 2 | 100% (10/10) | 90% (9/10) | 60% (6/10) | 60% (6/10) |
| Week 3 | 80% (8/10) | 90% (9/10) | 80% (8/10) | 80% (8/10) |
| Week 4 | 90% (9/10) | 90% (9/10) | 60% (6/10) | 70% (7/10) |
| Week 6 | 100% (10/10) | 100% (10/10) | 75% (6/8) | 80% (8/10) |
| Week 8 | 100% (9/9) | 90% (9/10) | 100% (9/9) | 80% (8/10) |
| Total | 95% (56/59) | 93% (55/59) | 79% (45/57) | 77% (46/60) |
|  | 94% (111/118) |  | 78% (91/117)* |  |

\*Significantly lower incidence compared to the endometrial tissue animals with and without estrogen. Fisher's exact test and Chi-square test were performed for statistical analysis.

The average number of 2.5±1.4 attached peritoneal implants per animal (out of 10 implanted tissue pieces) was significantly higher (p<0.0001) in the endometrial tissue mice, compared to adipose tissue animals (1.6±1.5 attached peritoneal implants; Figure 2A). The estrogen-substituted animals showed a peritoneal implant take rate of 2.1±1.4 attached peritoneal implants (combined results of endometrial- and adipose tissue animals), which was not significantly different (p=0.84) from the average take rate of 2.1±1.6 attached peritoneal implants of the estrogen-depleted animals (combined results of endometrial- and adipose tissue animals). The number of attached peritoneal implants was equal at the different time points of follow-up (1, 2, 3, 4, 6, or 8 weeks) (p=0.18 estrogen-substituted endometrial tissue animals; p=0.63 estrogen-depleted endometrial tissue animals; p=0.21 estrogen-substituted adipose tissue animals; p=0.34 estrogen-depleted adipose tissue animals), indicating that there was no significant regression in in number of attached peritoneal implants over time. Moreover, red, brown, yellow and white colored attached peritoneal implants were observed in animals after induction of both menstrual endometrium and adipose tissue, see Figure 3.

**Figure 2:**
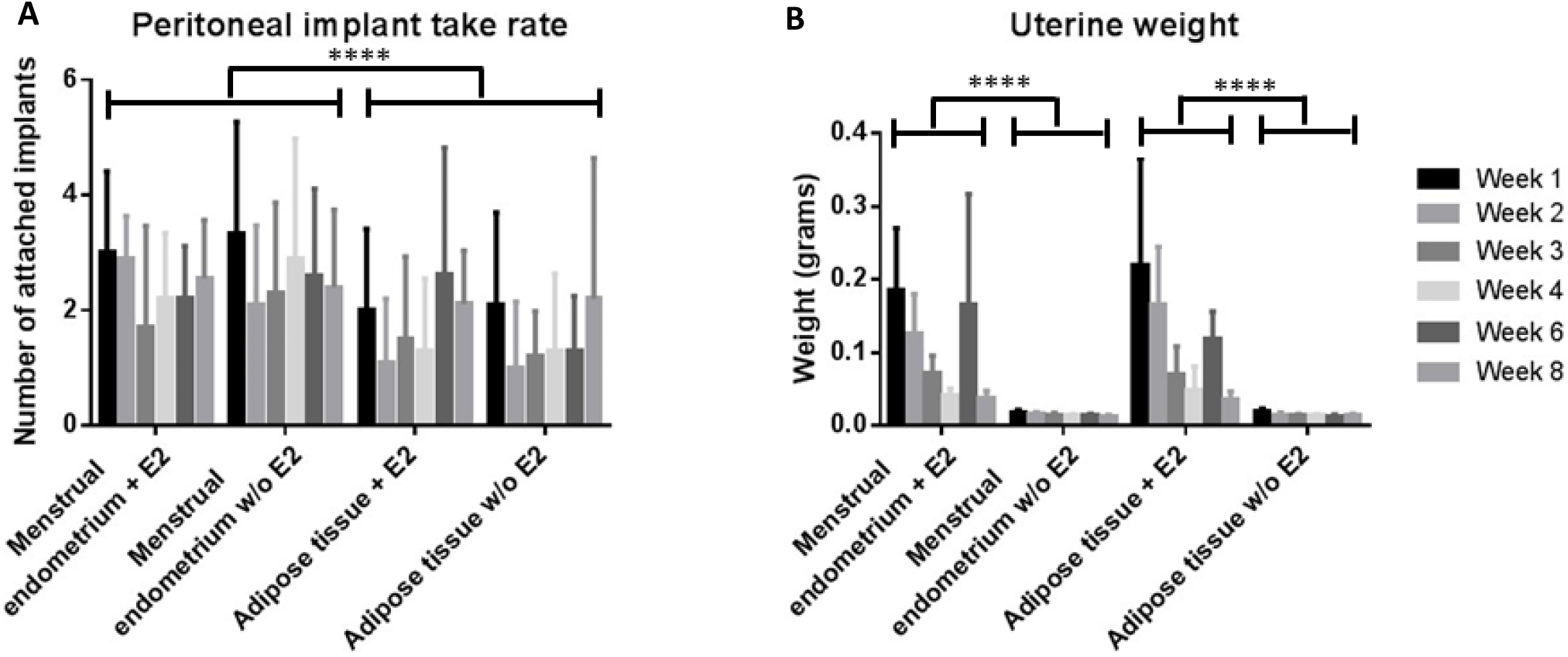
Number of attached implants and uterine weight of the recipient animals. **A:** The average number of attached peritoneal implants was significantly higher (p<0.0001) in the endometrial tissue mice, compared to adipose tissue animals. Estrogen supplementation and time did not show to affect the number of attached peritoneal implants. **B:** A significantly higher average uterine weight was observed in the estrogen-substituted animals, compared to the estrogen-depleted animals. Furthermore, the estrogen-supplemented animals showed an estrogen-related reduction of uterine weight, which could be undone by replacement of the estrogen capsule four weeks after placement of the initial capsule, as shown by the increase in uterine weight at 6 weeks. A two-way ANOVA was performed. Data are depicted as mean + standard deviation.

**Figure 3:**
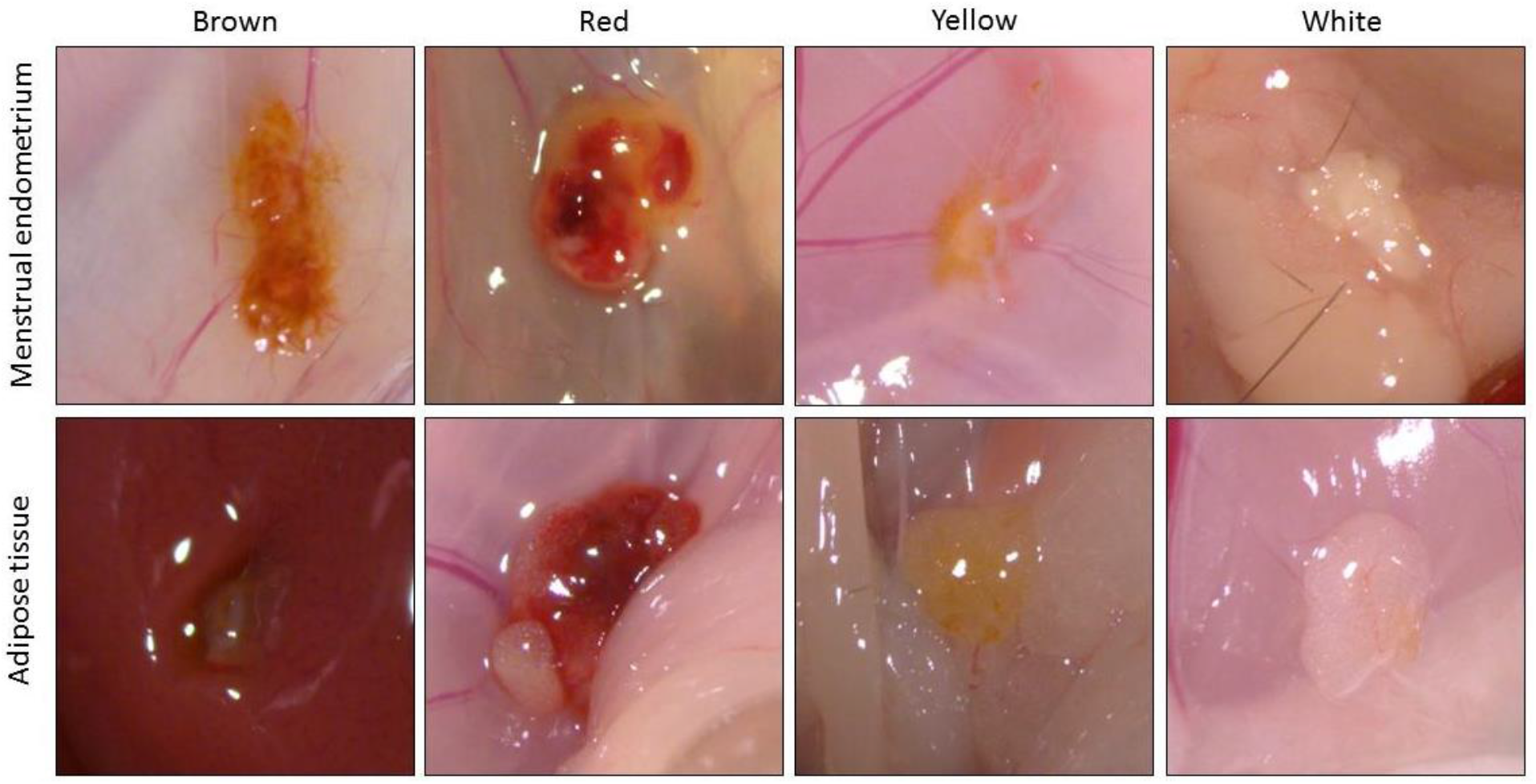
Macroscopic appearance of attached peritoneal implants at sacrifice.

The retrieved peritoneal implants showed an average length of 1.91±0.87 mm. The length of the attached peritoneal implants was equal between the different groups (p=0.07) and time points (p=0.56).

#### 3.2.2. Uterine weight

The average uterine weight of the estrogen-substituted animals was 109±96 mg, which was significantly higher (p<0.0001) compared to the estrogen-depleted animals, in which an average uterine weight of 15±3 mg could be observed (Figure 2B). Furthermore, an estrogen-related reduction of uterine weight was observed over the first four weeks in the estrogen-supplemented animals (p<0.0001; Figure 2B), due to a known decrease in estrogen release over time by the capsule. This effect could be undone by replacement of the estrogen capsule four weeks after placement of the initial capsule, as shown by the increase in uterine weight at week 6 compared to week 4 (p<0.01).

#### 3.2.3. Hematoxylin and eosin staining

One attached peritoneal implant from each mouse was analyzed using H&E staining. 33 animals were excluded from analysis, due to a lack of attached lesions. Furthermore, 18 animals were excluded due to technical artifacts, such as lesions that were too small for embedding or no observation of endometrial-like or adipose tissue in the sections.

H&E staining of the included attached implants revealed that none of the implants showed the typical endometriotic structure of glands and stroma. The majority of cells in the attached peritoneal implants were mononuclear immune cells, such as lymphocytes and macrophages, in addition to multi-nucleated giant cells. Furthermore, poly-nuclear immune cells, such as neutrophils and eosinophils, could be observed in the attached peritoneal implants as well. We also showed the presence of fibrotic tissue in the attached implants. In some of the endometrial tissue animals, stromal cells were observed and found to be decidualized cells, while in the attached peritoneal implants of the adipose tissue animals, the presence of adipose cells could be observed (Figure 4).

**Figure 4:**
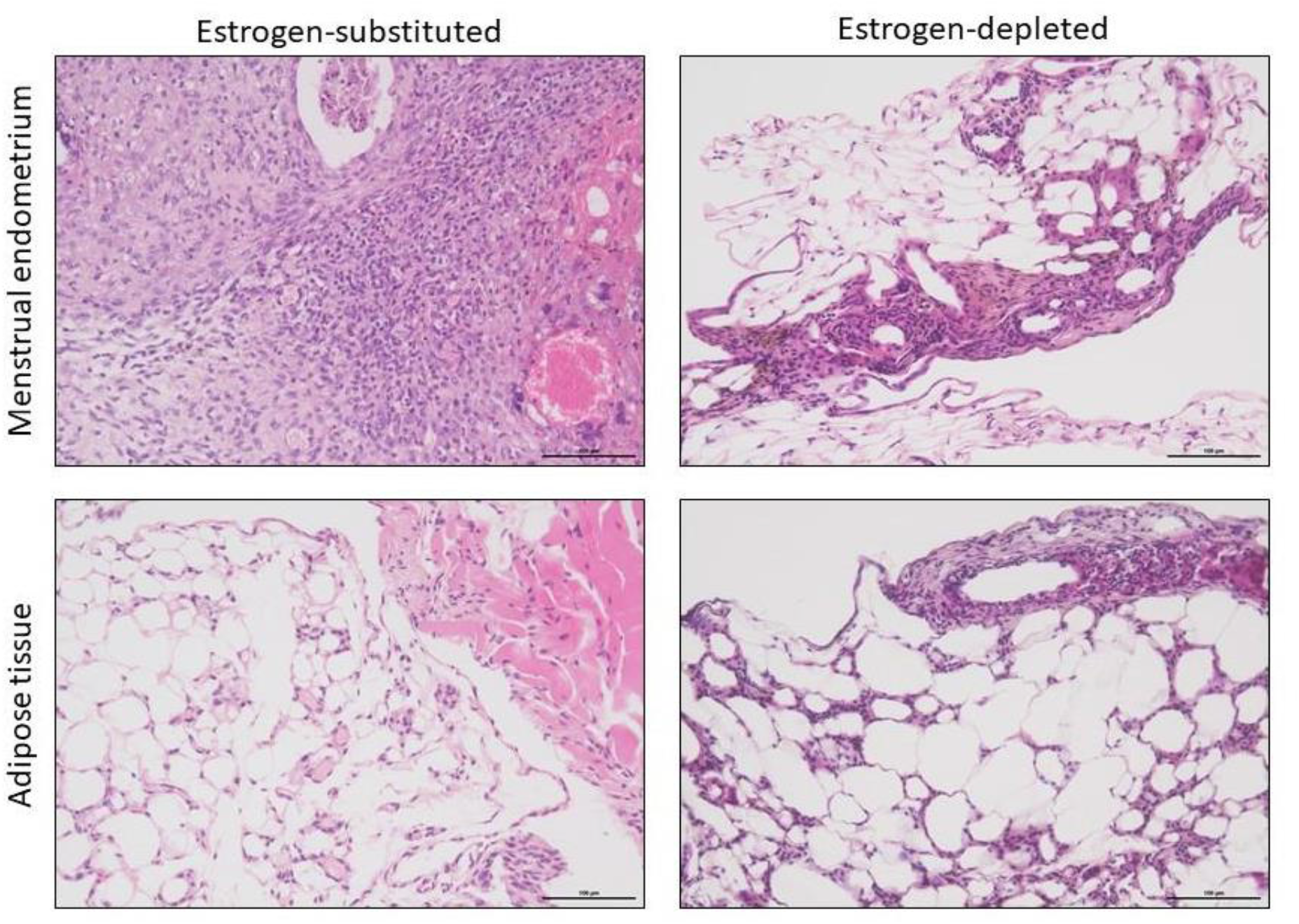
Microscopic visualization of the attached peritoneal implants using H&E staining. Attached peritoneal implants show the presence of stromal cells (menstrual endometrium) or adipose cells (adipose tissue) with different types of immune cells and hemosiderin (picture of menstrual endometrium, estrogen-depleted attached peritoneal implant). Scale bar: 100µm

Furthermore, the presence of hemosiderin in the tissues was analyzed and shown to be significantly higher (p<0.0001) in the implants induced with menstrual endometrium compared to the peritoneal implants of the adipose tissue animals, regardless of estrogen supplementation (endometrial tissue: p=0.83; adipose tissue: p=0.37). Additionally, the number of attached peritoneal implants showing hemosiderin increased over time in the endometrial tissue animals, with a significantly higher number of attached peritoneal implants showing hemosiderin from week 2 (estrogen-depleted animals) or week 3 (estrogen-substituted animals) onwards compared to week 1 (Table 3).

**Table 3:** Presence of hemosiderin in HE staining in attached peritoneal implants of recipient mice.

|  | Endometrial tissue + E<sub>2</sub> | Endometrial tissue – E<sub>2</sub> | Adipose tissue + E<sub>2</sub> | Adipose tissue – E<sub>2</sub> |
| --- | --- | --- | --- | --- |
| Week 1 | 20% (2/10) | 11.1% (1/9) | 0% (0/8) | 37.5% (3/8) |
| Week 2 | 50% (5/10) | 75% (6/8)<sup>a</sup> | 50% (3/6) | 75% (3/4) |
| Week 3 | 85.7% (6/7)<sup>a</sup> | 87.5% (7/8)<sup>b</sup> | 16.7% (1/6) | 50% (4/8) |
| Week 4 | 87.5% (7/8)<sup>a</sup> | 62.5% (5/8)<sup>a</sup> | 66.7% (4/6)<sup>a</sup> | 71.4% (5/7) |
| Week 6 | 87.5% (7/8)<sup>a</sup> | 75% (6/8)<sup>a</sup> | 66.7% (4/6)<sup>a</sup> | 37.5% (3/8) |
| Week 8 | 100% (8/8)<sup>b, c</sup> | 88.9% (8/9)<sup>b</sup> | 11.1% (1/9) | 0% (0/7)<sup>c, d</sup> |
| Total | 69% (35/51) | 66% (33/50) | 32% (13/41) | 43% (18/42) |
|  | 67% (68/101) |  | 37% (31/83)* |  |

<sup>a</sup>: $p<0.05$ compared to week 1
<sup>c</sup>: $p<0.05$ compared to week 2
<sup>b</sup>: $p<0.01$ compared to week 1
<sup>d</sup>: $p<0.05$ compared to week 4
\*Significantly lower incidence compared to the endometrial tissue animals with and without estrogen. Fisher's exact test and Chi-square test were performed for statistical analysis.

#### 3.2.4. Immunohistochemistry

One attached peritoneal implant from each mouse was analyzed using immunohistochemistry. 33 animals were excluded from immunohistochemical analysis, due to a lack of attached lesions. Furthermore, 18 animals were excluded for the PR and vimentin staining, 19 animals for the PGP9.5 and ERα staining, 23 animals for cytokeratin staining, 24 animals for Ki67 staining and 29 animals for podoplanin, due to technical artifacts, such as lesions that were too small for embedding, or no detection of endometrial-like or adipose tissue in the attached tissue after H&E staining.

All attached peritoneal implants stained positive for vimentin and 66% of the implants showed cytokeratin positive cells, without a difference between endometrial tissue and adipose tissue animals (p=0.53), nor between estrogen-supplemented and estrogen-depleted animals (p=0.08). Nevertheless, the cytokeratin positive cells appeared to be mesothelial cells, and were overgrowing the attached implants, rather than endometrial epithelial cells from the implanted tissue (Figure 5).

**Figure 5:**
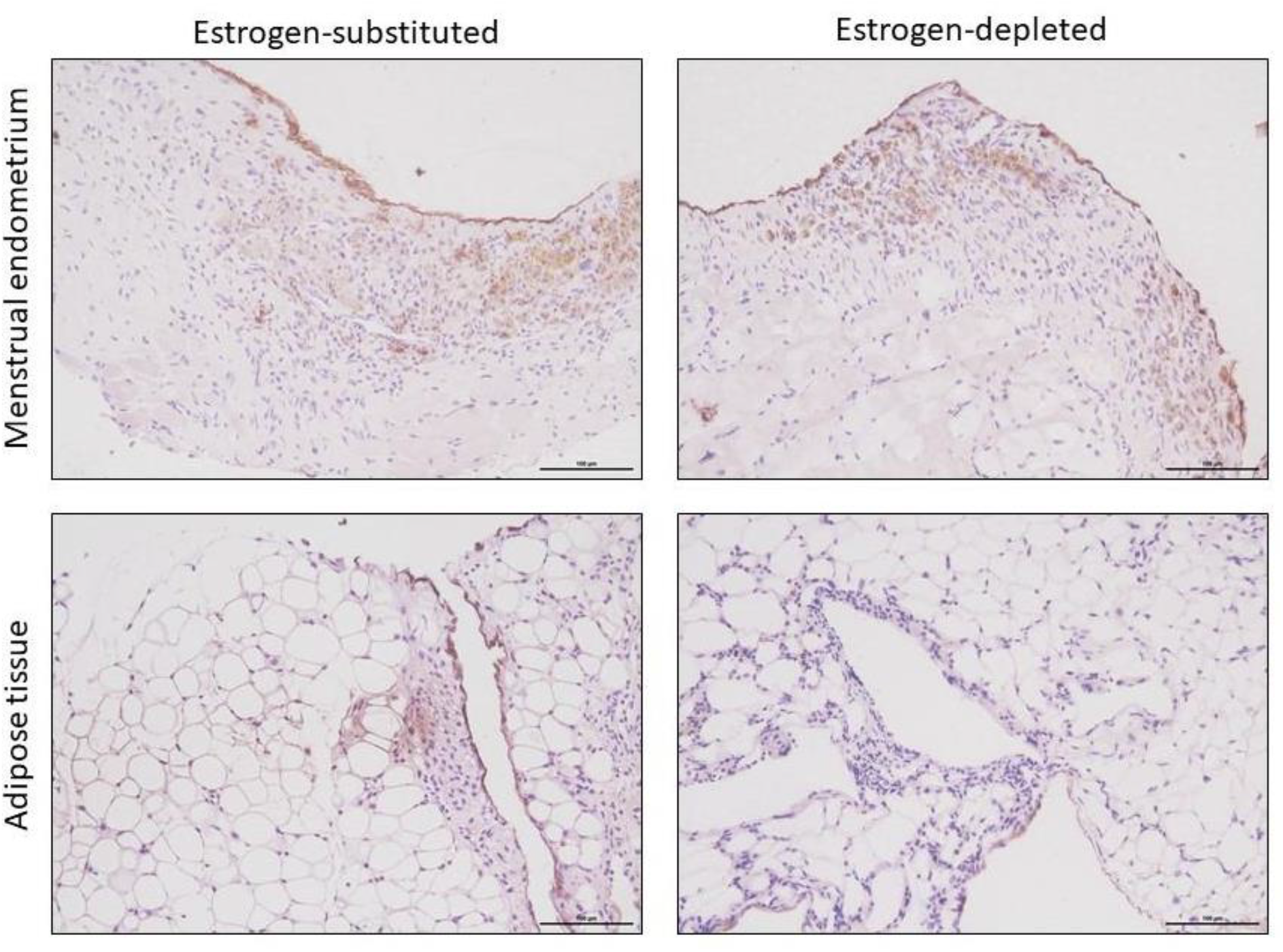
Immunohistochemical staining with a pan-cytokeratin marker. Positive staining for pan-cytokeratin can be observed as a single layer on top of the different attached peritoneal implants, indicating that these pan-cytokeratin positive cells are mesothelial cells that are overgrowing these implants. Scale bar: 100µm

Positive staining for the estrogen receptor alpha (ERα) and the progesterone receptor (PR) could be detected in all attached peritoneal implants. For PR, no significant difference in Allred scoring could be observed between endometrial tissue and adipose tissue animals, independent of estrogen substitution (p=0.94) and time point of sacrifice (p=0.27) (Supplementary Table 2). With respect to ERα staining, significant differences in Allred scoring could be observed (Table 4). Attached peritoneal implants from animals sacrificed one or two weeks after tissue implantation showed a significantly higher Allred score compared to the mice sacrificed eight weeks after tissue implantation (p<0.05). Additionally, the Allred score of the estrogen-depleted adipose tissue animals was significantly higher at week one and week three compared to week eight (p<0.05). Also, the Allred score of the estrogen-supplemented endometrial tissue mice was significantly higher at week four compared to week eight (p<0.05)

**Table 4:** Allred score after ERα staining in attached peritoneal implants of recipient mice.

|  | Endometrial<br>tissue + E<sub>2</sub> | Endometrial<br>tissue – E<sub>2</sub> | Adipose<br>tissue + E<sub>2</sub> | Adipose<br>tissue – E<sub>2</sub> |
| --- | --- | --- | --- | --- |
| Week 1 | 7.4±0.7 | 7.7±0.5 | 7.8±0.5 | 7.7±0.7 |
| Week 2 | 7.5±0.7 | 8.0±0.0 | 7.5±0.8 | 7.7±0.5 |
| Week 3 | 7.6±0.5 | 7.9±0.4 | 6.6±1.3 | 7.6±0.7 |
| Week 4 | 7.9±0.3 | 6.6±1.5 | 7.2±1.3 | 7.4±0.8 |
| Week 6 | 6.6±1.1 | 7.0±1.7 | 7.3±0.8 | 7.3±0.9 |
| Week 8<sup>a</sup> | 6.4±1.6<sup>b</sup> | 7.1±1.4 | 7.8±0.7 | 6.1±1.7<sup>c</sup> |
| Total | 7.3±1.0 | 7.4±1.1 | 7.4±1.0 | 7.3±1.1 |
|  | 7.3±1.1 |  | 7.3±1.0 |  |

<sup>a</sup>: $p < 0.05$ compared to week 1 and week 2
<sup>c</sup>: $p < 0.05$ compared to week 1 and week 3
<sup>b</sup>: $p < 0.05$ compared to week 4
A two-way ANOVA was performed for statistical analysis. Data are depicted as mean ± standard deviation.

Pgp9.5 staining was performed to identify nerve fibers in the attached peritoneal implants (Figure 6). This staining revealed that an equal number of nerve fibers per mm^2^ could be observed in the attached peritoneal implants of the endometrial tissue (2.4±7.3 nerve fibers/mm^2^) and adipose tissue animals (3.6±11.6 nerve fibers/mm^2^), independent of estrogen-substitution (p=0.59, estrogen substituted mice: 2.3±6.9 nerve fibers/mm^2^; estrogen-depleted animals: 3.7±11.6 nerve fibers/mm^2^; see Supplementary table 3).

**Figure 6:**
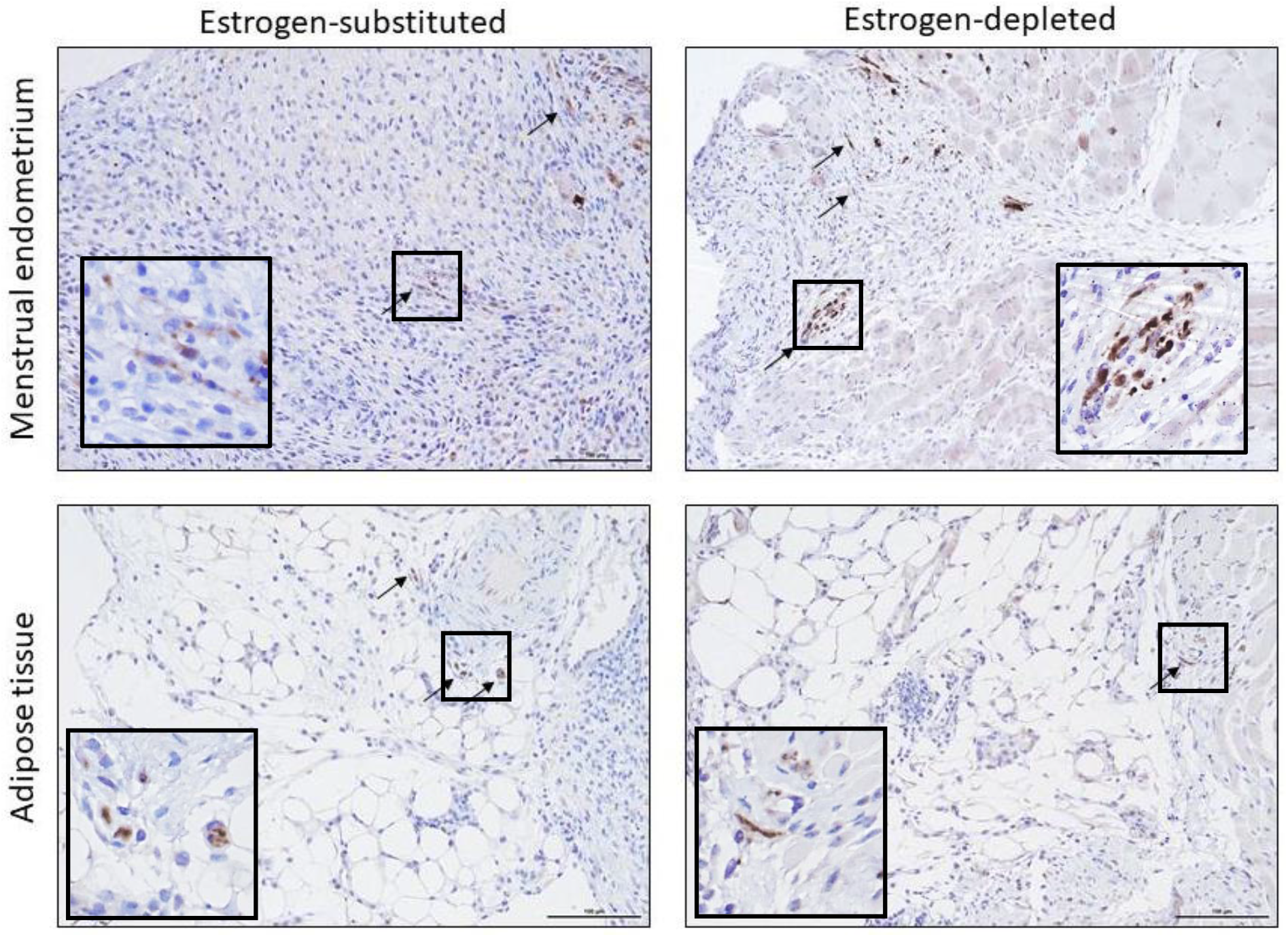
Immunohistochemical staining with a PGP9.5. Nerve fibers could be detected in all attached implants, irrespective of type of implanted tissue or hormonal status. Scale bar: 100µm

To investigate cell proliferation, Ki67 staining was used as marker and showed that the number of proliferative cells was significantly higher at week 1 after tissue implantation, compared to the other time points (p<0.001 vs. week 2; p<0.0001 vs. week 3, week 4, week 6 and week 8, Figure 7). There was no difference in proliferation index (p=0.14) between the endometrial tissue (estrogen-substituted: 15.3±9.5%; estrogen-depleted: 12.3±9.4%) and adipose tissue animals with and without estrogen-supplementation (estrogen-substituted: 17.3±13.9%; estrogen-depleted:14.7±12.6%).

**Figure 7:**
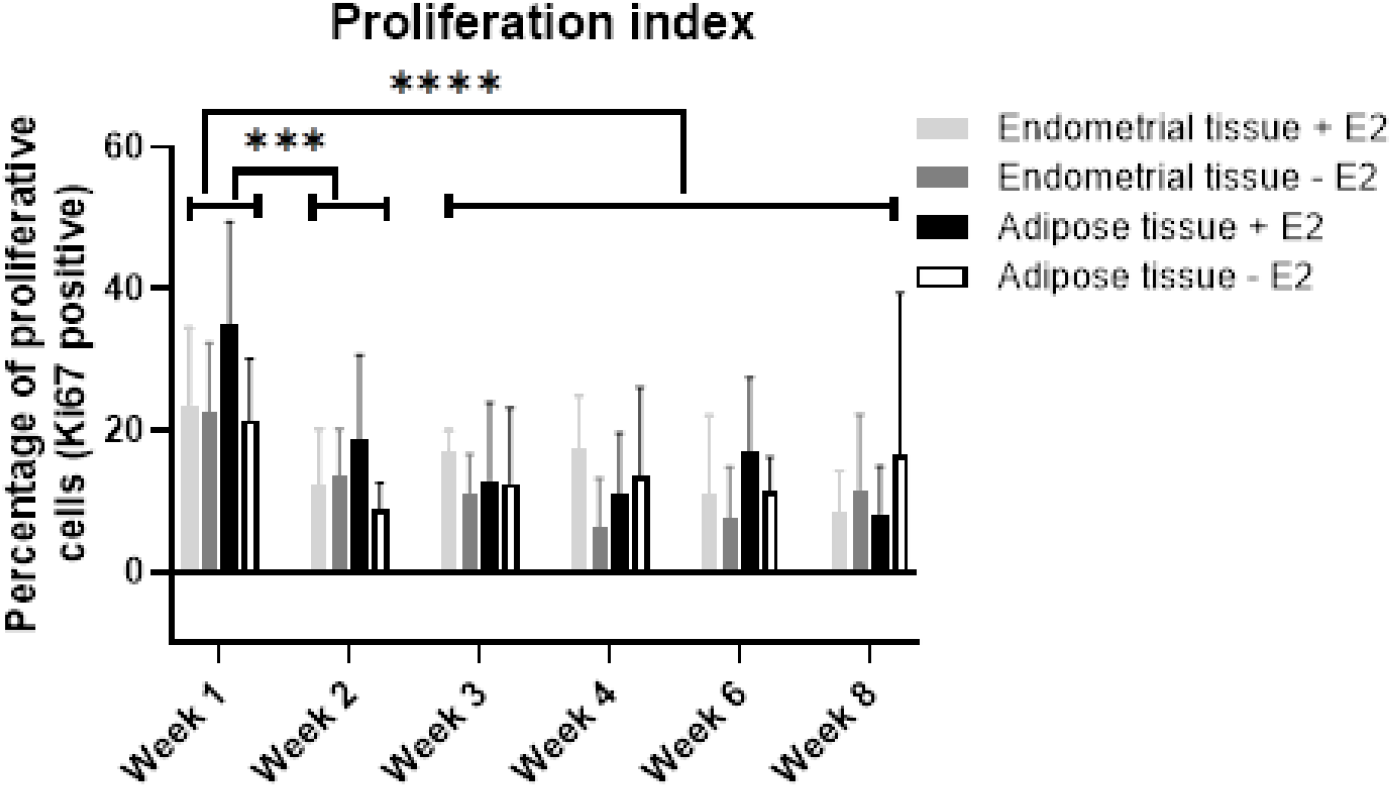
Proliferation index after Ki67 staining in attached peritoneal implants of recipient mice. A significantly higher percentage of proliferative cells could be detected in attached peritoneal implants at week 1 compared to all other weeks (p<0.001 compared to week 2; p<0.0001 compared to weeks 3, 4, 6, and 8). There was no difference in percentage of proliferative cells between endometrial or adipose tissue, nor between estrogen-substituted and estrogen-depleted. A two-way ANOVA was performed for statistical analysis. Data are depicted as mean + standard deviation.

Immunohistochemical staining with an antibody against podoplanin revealed an equal number of lymph vessels in the attached peritoneal implants of endometrial tissue and adipose tissue animals p=0.09), independent of estrogen substitution and time point of sacrifice (p=0.46) (see Supplementary Table 4).

#### 3.2.5. Behavioral tests

For the endometrial tissue group, only animals with at least one retrieved peritoneal implant were included in the analyses, while for the adipose tissue group, all animals were included.

We observed that 51% (118/230) of the recipient mice were in the estrus phase of the cycle during these first series of behavioral tests, 22% (51/230) of the animals were in the metestrus phase, 14% (32/230) were in diestrus and 13% (29/230) showed to be in the proestrus phase of the estrous cycle.

##### Thermal sensitivity

Behavioral tests measuring thermal sensitivity detected that estrogen-substituted animals were less sensitive to thermal stimuli compared to the estrogen-depleted mice (the results of both endometrial- and adipose tissue animals have been combined for all estrogen-related data).

In the acetone test, which is a measure for cold sensitivity, the estrogen-substituted animals showed a stronger reduction in total reaction time (p<0.05, Figure 8A) and times they exhibited abdominal grooming (p<0.05, Figure 8B) after surgery compared to before, than the estrogen-depleted animals. In addition, the number of times that the animals groomed their abdomen decreased significantly over the weeks (p<0.05, Figure 8B) and was significantly lower after tissue implantation compared to before (p<0.0001, Figure 8B).

**Figure 8:**
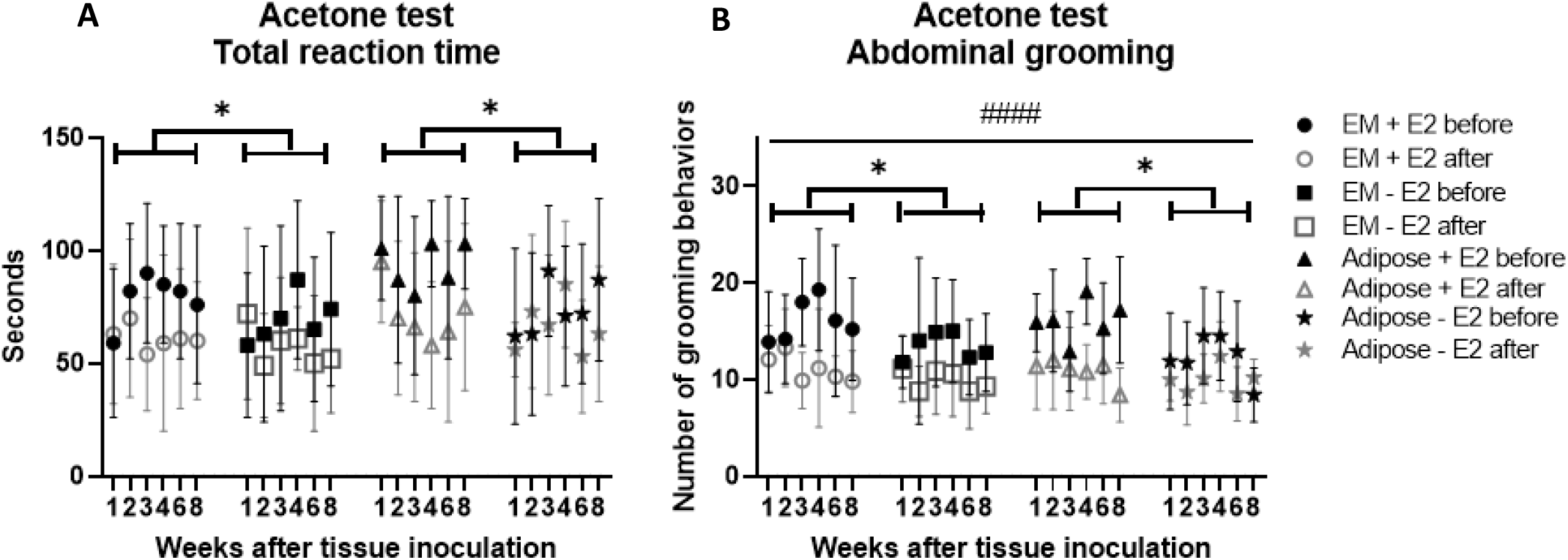
Total reaction time and abdominal grooming behavior in acetone test. **A:** Total reaction time in seconds after application of acetone onto the lower abdomen. A significant decrease in total reaction time could be observed after tissue implantation compared to before in the estrogen-supplemented animals (p<0.05 for reduction of total reaction time compared to estrogen-depleted animals). **B:** Total number of abdominal grooming behaviors exhibited by the animals after applying acetone onto the lower abdomen before and after tissue implantation. A significant reduction of total number of abdominal grooming behaviors after tissue implantation compared to before (p<0.0001) and a significant reduction of total abdominal grooming behaviors of estrogen-supplemented animals compared to estrogen-depleted animals after surgery (p<0.05) could be observed. Linear models were applied for statistical analysis. Data are depicted as mean + standard deviation.

In the hot water test, estrogen-substituted mice showed an increased latency to first response at week four (p<0.05, Figure 9A) when compared to estrogen-depleted animals. Moreover, the number of times the animals were showing abdominal grooming was significantly different across weeks (p<0.01, Figure 9B) and was significantly lower after tissue implantation compared to before (p<0.01, Figure 9B)

**Figure 9:**
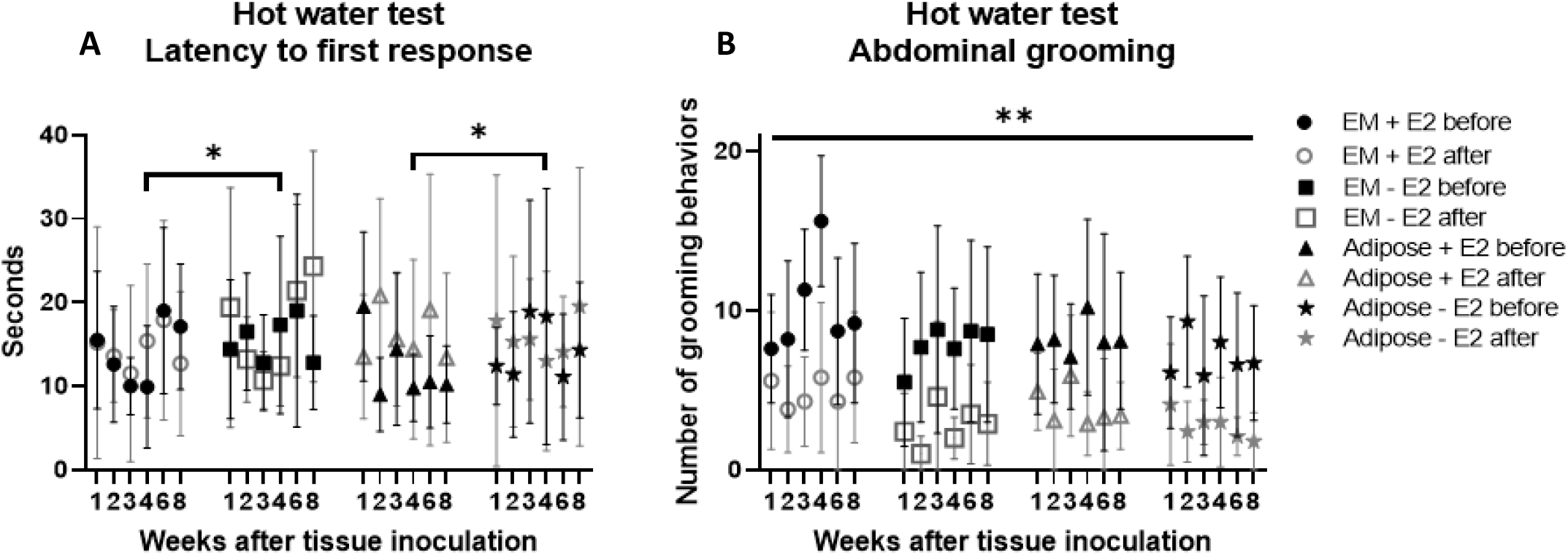
Latency to first response and abdominal grooming behavior in hot water test. **A:** Latency to first response in seconds after application of hot water (PEG) onto the lower abdomen. A significant difference in the latency to first response before surgery versus after surgery could be observed at 4 weeks in the estrogen-supplemented animals compared to the estrogen-depleted ones (p<0.05). **B:** Total number of abdominal grooming behaviors exhibited by the animals after applying hot water onto the lower abdomen before and after tissue implantation. The number of times that animals groomed their abdomen decreased significantly over the weeks (p<0.01) and a significant reduction of total number of abdominal grooming behaviors after tissue implantation compared could be observed in all groups/timepoints to before (p<0.01). Linear models were applied for statistical analysis. Data are depicted as mean + standard deviation.

In the hargreaves test, a prolonged withdrawal threshold in seconds could be observed after surgery versus before at week 3 (p<0.001) and week 6 (p<0.05) in the estrogen-substituted animals (Figure 10), when compared to the estrogen-depleted animals.

**Figure 10:**
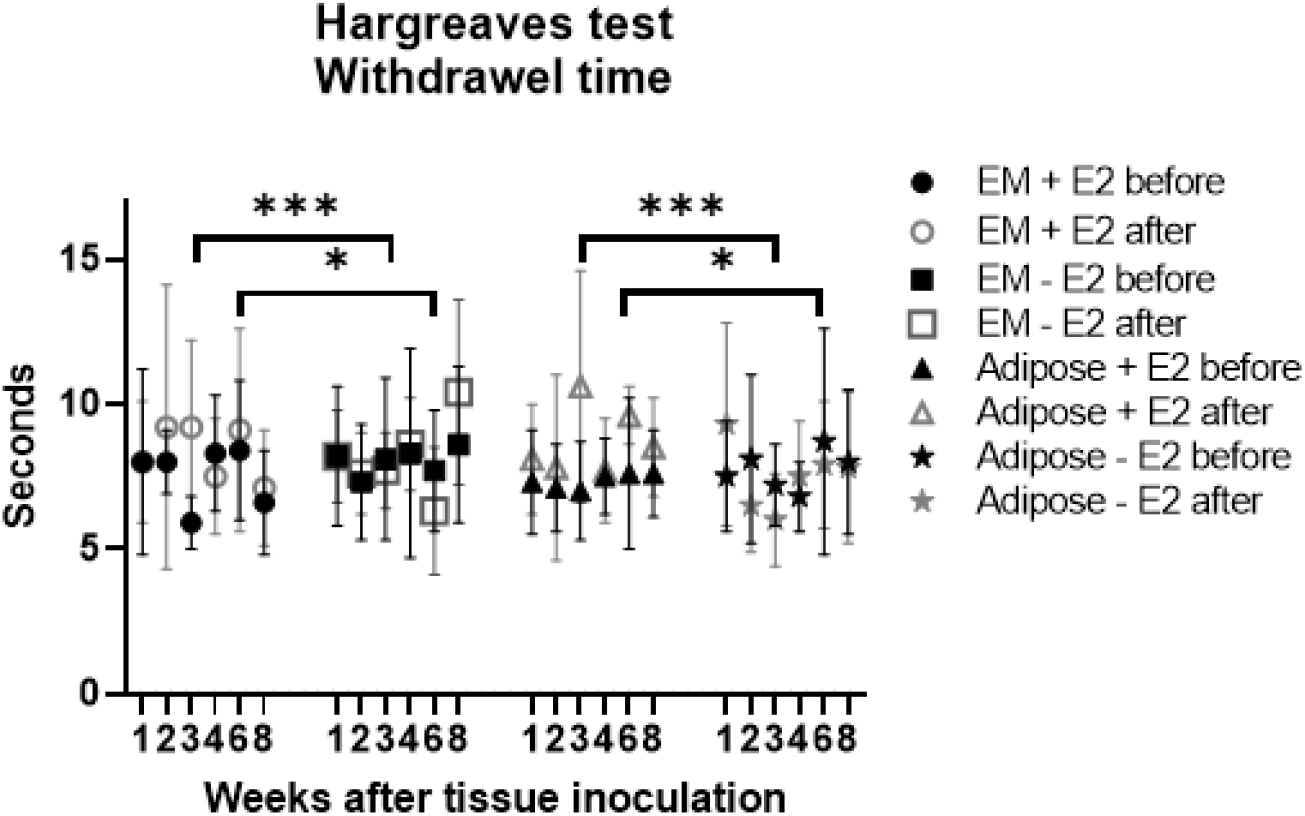
Withdrawel threshold (s) in Hargreaves test. Withdrawel threshold in seconds using the Hargreaves test before and after tissue inoculation in the recipient animals. A significant difference could be observed in the withdrawel response before versus after surgery in estrogen-supplemented animals compared to the estrogen-depleted animals at week 3 (p<0.001) and week 6 (p<0.05). Linear models were applied for statistical analysis. Data are depicted as mean + standard deviation.

##### Mechanical sensitivity

In the Von Frey hair test, which measures mechanical sensitivity, no significant difference in withdrawal threshold could be observed between the animals in which menstrual endometrium was inoculated compared to the animals which received adipose tissue, neither between the estrogen-substituted animals compared to the estrogen-depleted ones (p=0.30), nor could a difference in withdrawal threshold be observed over time (p=0.60) (see Supplementary Table 5).

##### Psycho-physiological tests

The results of the open field test indicated that the distance walked in the periphery was significantly lower after implantation of both menstrual endometrium and adipose tissue, compared to before (p<0.0001; Figure 11), although this distance walked increased after tissue implantation over the weeks in all groups (p<0.05). Moreover, the open field test indicated that estrogen did not alter voluntary activity of the animals after tissue implantation, when comparing estrogen-substituted animals with the estrogen-depleted mice.

**Figure 11:**
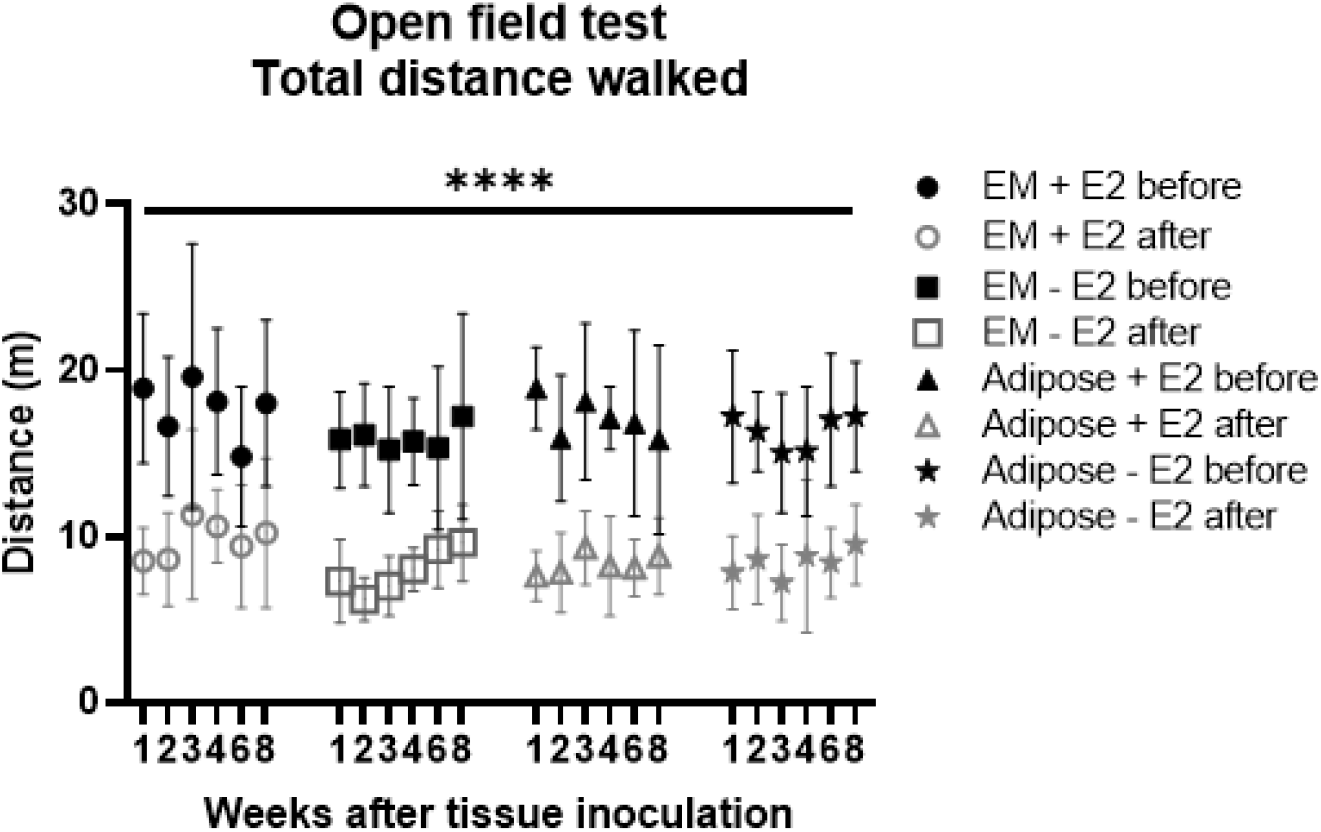
Total distance (m) walked in the periphery of the open field. A significant decrease in total distance walked in the periphery could be observed in all groups and timepoints after surgery compared to before surgery (p<0.0001). Linear models were applied for statistical analysis. Data are depicted as mean + standard deviation.

The results of the marble burying test (Table 12) showed that the estrogen-substituted animals buried less marbles after surgery than before compared to the estrogen-depleted mice (p<0.05). The total number of marbles buried by the animals changed over the weeks (p<0.05), with a trend towards the animals burying more marbles after surgery compared to before during the first 3 weeks and burying less marbles after surgery compared to before starting from week 4 onwards, with a significant difference in week 8 (p<0.01). Additionally, the number of buried marbles increased over time in each experimental session (p<0.05).

**Table 12:** Average number of buried marbles of the recipient animals before and after tissue inoculation using the Marble burying test.

|  |  | Endometrial tissue + E2<sup>a</sup> |  | Endometrial tissue – E2 |  | Adipose tissue + E2<sup>a</sup> |  | Adipose tissue – E2 |  |
| --- | --- | --- | --- | --- | --- | --- | --- | --- | --- |
|  |  | Before | After | Before | After | Before | After | Before | After |
| Week 1 | 5 min | 0.0±0.0 | 1.7±2.1 | 0.6±1.1 | 1.8±2.7 | 1.8±2.1 | 2.2±3.6 | 1.5±4.7 | 2.1±4.1 |
|  | 10 min | 1.2±2.4 | 2.9±3.3 | 3.0±5.3 | 4.1±4.9 | 3.4±3.9 | 5.5±5.1 | 2.7±3.8 | 4.3±4.1 |
|  | 15 min | 3.0±3.1 | 4.5±5.0 | 4.3±5.1 | 5.1±4.5 | 5.1±5.1 | 6.4±5.0 | 4.7±4.5 | 6.4±4.3 |
|  | 20 min | 5.7±4.4 | 6.6±5.1 | 5.8±4.9 | 6.1±4.1 | 7.5±6.1 | 8.1±5.0 | 6.6±4.8 | 7.7±4.3 |
|  | 25 min | 7.2±4.5 | 8.2±5.5 | 8.1±5.6 | 8.0±5.0 | 7.7±6.4 | 8.3±5.5 | 7.4±3.9 | 9.3±4.5 |
|  | 30 min | 8.2±4.3 | 8.3±5.8 | 8.3±5.5 | 9.6±4.3 | 8.3±6.0 | 10.5±4.7 | 10.5±3.0 | 10.4±5.2 |
| Week 2 | 5 min | 0.5±1.6 | 1.3±1.4 | 0.5±0.8 | 1.5±1.4 | 1.3±1.4 | 0.9±1.5 | 0.2±0.7 | 0.9±1.1 |
|  | 10 min | 1.1±2.5 | 2.8±2.9 | 2.4±3.0 | 2.9±2.5 | 3.1±3.3 | 3.8±4.5 | 0.8±1.1 | 2.2±2.9 |
|  | 15 min | 2.8±4.3 | 4.2±4.2 | 5.1±4.0 | 4.6±3.5 | 5.7±4.2 | 4.7±4.5 | 2.7±4.1 | 4.4±4.1 |
|  | 20 min | 3.7±4.7 | 6.1±5.4 | 6.9±4.1 | 6.0±4.4 | 7.3±4.9 | 6.4±4.8 | 3.7±4.2 | 6.0±4.5 |
|  | 25 min | 4.4±5.4 | 6.7±5.5 | 8.8±3.8 | 9.4±3.7 | 10.5±5.8 | 6.2±4.9 | 5.4±4.9 | 6.6±3.8 |
|  | 30 min | 4.4±5.8 | 6.7±4.5 | 9.9±3.6 | 9.8±5.1 | 10.3±5.9 | 7.4±4.4 | 7.6±4.2 | 9.2±3.2 |
| Week 3 | 5 min | 0.6±1.4 | 0.4±0.5 | 0.8±2.1 | 0.8±1.8 | 0.8±1.5 | 1.8±2.0 | 0.0±0.0 | 0.6±1.1 |
|  | 10 min | 1.1±1.8 | 1.4±1.3 | 1.9±3.4 | 2.9±4.8 | 3.4±4.9 | 4.2±4.1 | 1.1±2.4 | 1.6±1.8 |
|  | 15 min | 2.3±3.1 | 2.8±2.1 | 3.0±3.3 | 3.6±4.7 | 5.8±4.7 | 7.2±5.6 | 2.9±4.1 | 2.7±2.5 |
|  | 20 min | 3.5±4.1 | 4.6±3.3 | 4.9±4.8 | 5.9±4.8 | 7.2±5.2 | 8.6±5.7 | 3.6±4.5 | 4.9±3.3 |
|  | 25 min | 5.3±6.1 | 5.0±3.7 | 6.9±6.1 | 7.5±4.4 | 10.0±5.8 | 9.5±4.8 | 5.2±4.6 | 7.2±3.4 |
|  | 30 min | 5.0±5.0 | 6.5±3.3 | 8.5±5.5 | 8.6±4.1 | 9.8±5.1 | 9.9±5.0 | 6.6±4.8 | 5.7±2.9 |
| Week 4 | 5 min | 0.4±0.7 | 0.6±1.0 | 1.0±2.6 | 2.7±2.2 | 0.4±0.8 | 0.4±1.0 | 0.9±1.6 | 0.2±0.4 |
|  | 10 min | 2.0±2.6 | 2.1±3.2 | 2.7±3.8 | 5.0±4.0 | 1.6±2.4 | 1.7±2.0 | 3.2±4.9 | 1.4±2.2 |
|  | 15 min | 3.7±3.8 | 3.6±4.2 | 5.8±5.6 | 7.0±5.2 | 3.2±3.9 | 2.4±2.6 | 6.0±5.9 | 2.7±3.2 |
|  | 20 min | 4.6±4.3 | 5.8±5.4 | 7.4±4.1 | 8.6±4.6 | 5.1±5.1 | 3.7±3.3 | 6.9±4.5 | 3.5±4.2 |
|  | 25 min | 6.4±4.2 | 5.8±4.4 | 9.1±4.5 | 8.9±4.6 | 6.6±4.6 | 4.9±3.9 | 9.2±5.5 | 3.7±3.8 |
|  | 30 min | 6.8±4.9 | 6.7±4.2 | 11.6±2.8 | 9.9±3.8 | 7.2±5.0 | 6.6±4.1 | 11.1±4.0 | 5.3±4.1 |
| Week 6 | 5 min | 1.5±2.8 | 0.9±1.3 | 0.5±0.9 | 1.0±1.0 | 0.9±1.1 | 0.9±2.1 | 0.6±1.3 | 2.0±2.0 |
|  | 10 min | 3.0±3.9 | 1.4±1.5 | 2.2±3.2 | 3.7±2.5 | 1.4±1.8 | 1.0±1.9 | 2.1±2.9 | 3.6±3.3 |
|  | 15 min | 5.5±5.4 | 3.3±2.2 | 4.1±3.9 | 7.0±4.2 | 2.6±3.5 | 3.0±4.0 | 5.8±3.6 | 6.1±4.4 |
|  | 20 min | 6.3±5.5 | 4.0±3.0 | 6.2±4.8 | 7.5±3.4 | 4.6±5.0 | 4.3±3.6 | 8.2±3.4 | 6.7±4.3 |
|  | 25 min | 6.2±5.2 | 4.6±3.1 | 8.5±5.5 | 6.9±2.3 | 4.9±5.0 | 6.5±4.5 | 9.7±3.8 | 8.2±3.9 |
|  | 30 min | 8.4±5.4 | 5.4±3.7 | 10.7±5.3 | 8.9±3.0 | 7.8±6.1 | 8.4±4.0 | 11.3±2.5 | 7.9±3.8 |
| Week 8<sup>b</sup> | 5 min | 3.0±4.4 | 0.8±1.7 | 0.1±0.4 | 0.9±2.1 | 0.8±1.1 | 0.3±0.5 | 0.4±0.7 | 0.2±0.4 |
|  | 10 min | 3.7±3.8 | 1.7±3.9 | 1.9±2.9 | 2.1±3.4 | 2.1±2.8 | 0.9±1.3 | 3.4±2.6 | 1.6±1.9 |
|  | 15 min | 8.0±4.4 | 3.2±3.9 | 2.0±3.2 | 4.6±4.7 | 3.4±4.8 | 2.3±2.6 | 5.4±2.8 | 4.1±2.8 |
|  | 20 min | 9.3±4.8 | 4.0±4.3 | 3.3±2.2 | 4.6±4.4 | 4.8±4.1 | 5.0±4.7 | 8.2±3.5 | 5.3±2.9 |
|  | 25 min | 9.9±4.0 | 4.8±4.0 | 5.5±4.4 | 5.3±4.8 | 6.2±4.9 | 7.0±4.0 | 8.1±1.9 | 8.0±3.0 |
|  | 30 min | 10.4±3.4 | 6.7±4.0 | 6.4±3.5 | 6.4±3.5 | 7.7±5.6 | 7.2±3.4 | 10.0±1.6 | 10.6±3.4 |

Total number of buried marbles significantly increased over 30 minutes in each session ( $p<0.05$ ), while total number of buried marbles changed over the weeks ( $p<0.05$ ). <sup>a</sup>: Number of buried marbles. $p<0.05$ compared to estrogen-depleted animals. <sup>b</sup>: Number of buried marbles. $p<0.01$ compared to before surgery. Linear models based on the longitudinal data were applied for statistical analysis.

## 4. DISCUSSION

Currently published mouse models of endometriosis use different methodologies regarding hormonal environment, method of induction, type of implanted tissue, and interval between induction and evaluation, resulting in contradictory research results^19, 25, 30, 33, 37, 68–71^. Therefore, we initiated this study in which we investigated lesion ontogenesis, endometriosis-related neurogenesis and endometriosis-associated pain in normal estrogenic and hypo-estrogenic conditions at several time points in a mouse model induced with menstrual endometrium.

In this study, we indicated that the appearance of the attached peritoneal implants changed most dynamically within the first two weeks after implantation, as shown by a higher number of proliferative cells in all attached implants, and an increasing number of hemosiderin-positive attached implants in the animals in which menstrual endometrium was implanted. Additionally, we showed that pain behavior did not only occur after attachment of endometrium-like tissue, but also after attachment of adipose tissue. These results highlight the need for adequate controls in animal studies for endometriosis, allowing to differentiate between endometriosis-associated pain and pain caused by other reasons. In our study, the observed pain behavior could also be chronic surgery-related pain and not endometriosis-associated pain.

### 4.1. Tissue attachment

In our study, we observed a peritoneal implant take rate of 25% (2.5±1.4 attached peritoneal implant out of 10 implanted tissue pieces), which was somewhat lower than the 50% observed in our previous pilot study using a similar model^24^. This inconsistency in results could possibly be explained by the learning curve related to laparoscopic surgery, since it has previously been shown that laparoscopy in mice is a complex technique, for which specific training and skills are needed, with a learning curve in both experienced and inexperienced surgeons^72^. An increasing surgical experience leads to a shorter duration of surgery, hereby causing less damage to the peritoneal wall, which could possibly negatively affect peritoneal implant take rate. Furthermore, this difference could be related to circulating estrogen and progesterone levels in the recipient animals, since natural cycling animals were used in the previous study, while ovariectomized and estrogen-substituted mice were used in this study. Previously, it has been shown that the estrogen serum levels are comparable between natural cycling animals and ovariectomized animals substituted with this type of silastic capsule^50^. Nevertheless, the possibility that the natural variation in circulating estrogen levels favors tissue attachment in endometriosis should not be neglected, although tissue attachment in natural cycling animals *versus* ovariectomized and estrogen-substituted animals has never been compared.

In other articles using an endometriosis mouse model injected with menstrual tissue, the actual peritoneal implant take rate, as defined in our study, was not reported, but it was mentioned that an average of 2.1 attached lesions could be retrieved^23^. This number is comparable to our study, in which an average of 2.5 implants could be retrieved in the animals implanted with menstrual endometrium.

Besides, a take rate of 30-60% could be observed in homologous mouse models, injected with uterine or endometrial tissue^19, 49, 73, 74^. The lower take rate of 25% observed in our study could be explained by increased apoptosis in the menstrual endometrium, compared to the uterine or endometrial tissue used for endometriosis induction before, as it has been shown that endometrial apoptosis is detectable within 4 hours after progesterone withdrawal in mice with a decidualized endometrium^75^.

### 4.2. Tissue ontogenesis

The attached peritoneal implants derived from menstrual endometrium did not show the typical endometriotic features, existing of endometrial glands and stroma. This was a surprising observation, since previous studies using this menstruating mouse model for inducing endometriosis demonstrated the presence of both stromal and epithelial cells in most of the attached lesions^23–25^. In mice, the process of decidualization is characterized by stromal cells differentiating into decidual cells, while the luminal epithelial cells undergo apoptosis^76^. Therefore, only decidualized stromal cells were implanted into the peritoneal cavity of the recipient mice in studies using the menstruating mouse model for endometriosis. The presence of luminal and glandular epithelium in the endometriotic lesions of the previous studies can be cause by mesenchymal-to-epithelial transition (MET), during which endometrial mesenchymal stem cells transform into luminal and glandular epithelium under influence of progesterone and estrogen^45, 77, 78^. This process is stimulated by estrogen and progesterone, as an increase in hormonal levels initiates co-localization of STAT3 (Signal transducer and activator of transcription 3) with MCL-1 (Induced myeloid leukemia cell differentiation protein), which has been shown to cause mesenchymal-to-epithelial transition^78^. In the current study, estrogen was only supplemented starting from the moment of tissue inoculation, while in previous studies, estrogen was already present in the animals at the moment that the decidualized and menstrual endometrium was inoculated^23–25^, and thus the window of MET induction might have been missed in our study.

Furthermore, we investigated for the first time the presence of hemosiderin in both menstrual endometrium and control (adipose) tissue, as previous investigators did not implant control tissue in their animals. We showed that hemosiderin was present in the attached peritoneal implants derived from both menstrual endometrium and adipose tissue, although the presence of hemosiderin was more common in the endometrial-like lesions. Previously, hemosiderin has also been found in endometriotic lesions in homologous and heterologous mouse models for endometriosis^23, 51, 70, 79^. The detection of hemosiderin in endometriotic lesions is caused by erythrocytes, which are present in the menstrual effluent^51^, while in the adipose tissue the presence of hemosiderin can be related to hemorrhage, caused by disruption of the blood vessel integrity after punching biopsies for inoculation into the recipient mice.

### 4.3. The role of estrogen in lesion ontogenesis

In this study, no effect of estrogen on tissue attachment, growth and maintenance could be detected in ovariectomized and estrogen-substituted mice, compared to the estrogen-depleted ones. This could be explained by the observation that estrogen increases glandular secretory activity, epithelial cell height, and epithelial cell proliferation, of which the latter two are regulated via ER-α^29, 36^. In our study, no epithelial cells could be observed in the attached peritoneal implants, and therefore, we were not able to find an estrogen-related difference in take rate, lesion size and - appearance.

Moreover, our results are comparable to investigators who also showed no effect of estrogen on take rate^33^, lesion size, and tissue pattern^27, 28^ in estrogen-substituted and natural cycling mouse models for endometriosis. However, our data are in contrast with results from other investigators observing an estrogen-related difference in take rate, lesion size, or appearance^27, 29, 30, 35, 36^, which could possibly be explained by differences in study set-up. Indeed, in our study, murine menstrual endometrium, existing of decidualized stromal cells, was induced, while uterine or endometrial tissue, existing of stromal and epithelial cells, has been used in other studies^27, 29, 30, 35, 36^. Another explanation for these conflicting results could be the difference in amount of supplemented estrogen, as the dose of administered estrogen was around 10-15 times higher in the other studies^29, 30, 36^ compared to ours, implying that the amount of substituted estrogen was not enough to induce a estrogen-related difference in lesion size nor appearance. We can conclude from these conflicting results that the role of estrogen in lesion development still is not clear.

### 4.4. The role of time in lesion ontogenesis

In this study, we showed that the variable “time since induction” (1, 2, 3, 4, 6 or 8 weeks) did not affect number nor size of attached peritoneal implants. The equal number of attached implants at all time points could be explained by the lack of iterative seeding of the menstrual endometrium, implying that an increase in take rate was not possible. Nevertheless, recent studies did observe an increase in the number of lesions over time^80, 81^. On the other hand, the number of attached peritoneal implants did not decrease over time either, which is not surprising, since it has been shown that attached lesions stabilize over time in rodent models for endometriosis^47, 82^, although others indicated a reduction in the number of lesions over time, depending on the type of tissue used for endometriosis induction^38^.

Furthermore, contrary to our findings, a positive correlation between time and lesion size has been observed in literature^83–85^. This discrepancy can be explained by the different type of tissue used to induce endometriosis, as menstrual endometrium was implanted in our study, while other investigators used uterine tissue, which can easily form cysts and hereby, increase lesion size^19, 69^.

Moreover, we showed that the proliferation index of the attached peritoneal implants was higher at week one, compared to the other time points. This suggests that early tissue attachment and generation require cell proliferation, while in later stadia of disease development, the cells only proliferate to maintain the status quo of the lesion. No difference in proliferation index could be observed between the sham and endometriosis animals, nor between the estrogen-substituted and estrogen-depleted mice. Contrary, Grümmer *et al.* showed that no difference in proliferation index could be observed over time in the estrogen-substituted animals, while the number of proliferating cells decreased dramatically over time in estrogen-depleted animals^30^. This inconsistency in results could possibly be explained by difference in the amount of administered estrogen, which was 0.2 µg per day in the study of Grümmer *et al.*^30^, and thus 10 times higher compared to our study, in which 0.7 µg estrogen was released over a 28-day period.

### 4.5. Neurogenesis

The results of our Pgp9.5 staining indicated that nerve fibers were present in the attached peritoneal implants from both menstrual endometrium and adipose tissue, without an increase in number of observed nerve fibers over time, or in relation to estrogen.

In endometriosis mouse models injected with menstrual endometrium, the presence of nerve fibers in the retrieved lesions has been shown before^41, 43^. Furthermore, innervation of the endometriotic lesions has been associated with the inflammatory milieu and with elevated levels of circulating estrogen^41, 43^.

Besides, the presence of nerve fibers in the adipose tissue could be related to adipose tissue initiated neurogenesis, since it has been shown that adipose tissue secretes neurotrophic factors and that adipose derived stem cells are able to initiate neural regeneration^86^. Moreover, it has been shown that white adipose tissue is innervated with sympathic and parasympathic nerve endings^87^.

### 4.6. Endometriosis-related behavior

In this study, we were not able to detect any endometriosis-associated pain behavior in our animals. These results are comparable to other investigators who indicated that no significant difference could be observed in thermal sensitivity between the endometriosis and sham animals^46, 88^, although others do document the presence of endometriosis-related hyperalgesia in mice^16, 44, 80, 81, 85, 89–92^. This discrepancy in results could be related to differences in study set-up, and more specifically in the choice of the control group. In our study, we implanted adipose tissue in the control animals and we showed that the mice implanted with menstrual endometrium and the control animals were comparable regarding tissue attachment, maintenance and nerve fiber innervation. Indeed, another study also showed limited differences in behavior between mice inoculated with endometrial tissue and mice implanted with adipose tissue^92^. Nevertheless, Castro et al^85^ indicated that endometriosis mice displayed increased sensitivity to evoked pain measurements, while sham animals, with attached adipose tissue implants, did not show such behavior. These contradictive results indicate that observed changes in pain related behavior are difficult to interpret and might or might not be endometriosis-specific. Several studies did show a positive effect on endometriosis-related behavior after treating endometriosis, with^46, 81, 88, 89, 91^ and without^16^ a reduction in lesion size. Treating the animals with known endometriosis medication might be a good strategy to identify if pain is endometriosis specific, even in cases where control tissue is not used or did not attach.

Estrogen substitution decreased thermal sensitivity and anxiety-like behavior in our study. It has been shown by some investigators that estrogen replacement in ovariectomized mice causes a decrease in anxiety-associated behavior and surgery-related hyperalgesia, although others demonstrated an estrogen-related increase in anxiety-related behavior^55–60^. To explain these discrepancies in behavior, it has been proposed that mice receiving estrogen experience more fear in situations that are already anxious to them^93^.

We showed that an increase in time between surgery and sacrifice (1, 2, 3, 4, 6, or 8 weeks) had a positive effect on thermal sensitivity and spontaneous behavior of the animals, as the mice became less sensitive to thermal stimuli, more active in the open field test and buried less marbles in the marble burying test over time. Nevertheless, other investigators showed contrary results and indicated no effect or a negative effect of time after surgery on thermal sensitivity and spontaneous behavior in either only endometriosis mice or in both endometriosis and sham animals^46, 81, 88, 92^. This discrepancy in results could be explained by using different animals for each time point in our study, while other investigators performed repetitive measurements on the same animals.

All animals in our study showed an increase in pain-related behavior after surgery compared to their situation before surgery. In the literature, post-operative chronic pain has been suggested to be dependent on the place and type of surgery^94^ and associated with altered behavior in different test settings^95^ ^96^. This may also explain the observed differences in behaviors before and after surgery in our study as each of the recipient animals underwent ovariectomy and laparoscopy before the second set of behavioral tests.

### 4.7. Future model improvement

The results of this study indicate that the methodology used to induce endometriosis in mice needs further optimization by firstly, increasing the implant take rate and secondly, ensuring that the retrieved implants contain both endometrial glands and stroma. We propose the following actions to achieve these goals.

The first possibility to increase the implant take rate would be by adjusting the duration of pneumoperitoneum, since it has been shown in an adhesion formation model that the number of adhesions increased with duration of pneumoperitoneum^97^. Secondly, the take rate could be enhanced by increasing the size of the inoculated tissue, since it has been suggested that larger tissue pieces are more difficult to clear by the immune system and therefore, more likely to adhere^98^. A third possibility would be to harvest the menstrual endometrium immediately after progesterone withdrawal in the donor mice, in order to avoid implantation of apoptotic tissue^75^.

Next, the presence of endometriotic structures, existing of glands and stroma, has to be observed in the attached tissue in order to state that the animals developed endometriosis. The first possibility to ensure the development of these endometriotic structures would be by implanting endometrial tissue from donor animals which did not show full bicornuate decidualization, suggesting that both glandular and luminal epithelial cells would still be present in the menstrual tissue of these donor mice, and that both stromal and epithelial cells would be inoculated in the recipient mice. A second option would be to use natural cycling recipient animals, similar to what has been described by Horne *et al.*^99^, since circulating estrogen and progesterone are suggested to regulate the process of mesenchymal-to-epithelial transition, during which endometrial mesenchymal stem cells transform into luminal and glandular epithelium^45, 77, 78^.

## 5. CONCLUSION

Taken together, we showed that both menstrual endometrium and adipose tissue can attach after laparoscopic inoculation in mice. The implant take rate was higher for menstrual tissue than for adipose tissue. After attachment, no differences could be observed regarding growth and maintenance between both tissue types. Implants induced with endometrial tissue did not fully phenocopy lesions seen in patients as they lacked a clear glandular and stromal tissue compartment. Moreover, no influence of estrogen could be observed regarding endometrial or adipose tissue attachment, growth, and maintenance. Additionally, pain behavior was not affected by the type of tissue used for induction, and the effect of estrogen replacement on behavior was limited. Overall, our study indicates that it is important to include a control group induced with control tissue in studies related to animal models in endometriosis, to ensure that outcomes are endometriosis specific.

## PERSONAL CONTRIBUTIONS

DP, DFO and AlV contributed to the design of the study, performing the experiments, analyzing the results, and writing and revising the article. AF, AV and ASVR were involved in designing the study, interpretation of the results and revising the article. PS, JV and TD contributed to the conception of the study, interpretation of the results and revising the paper.

## FUNDING

DP has received funding from Flanders Research Fund (FWO; number 11S6614N/11S6616N) to perform this research.

## CONFLICTS OF INTEREST

The authors AlV, AF, DFO, AV, A-SVR, PS, and JV declare no conflict of interest. Author DP reports only conflicts of interest outside the scope of the submitted manuscript. She has received a grant from Bayer Healthcare AG. Author TD reports only conflicts of interest outside the scope of the submitted manuscript. He has served as advisor for Bayer Pharma, Proteomika, Pharmaplex, Astellas, Roche Diagnostics, Actavis, has received grants from Ferring, Merck Serono, MSD, Besins, Pharmaplex, and has received travel support from Ferring, Merck Serono and MSD. From October 1st, 2015, Thomas D’Hooghe is Vice-President and Head of Global Medical Affairs Infertility for the Multinational Pharmaceutical company Merck Serono (Darmstadt, Germany) and continues on a part time basis his academic appointment as Professor of Reproductive Medicine at the University of Leuven (KU Leuven) in Belgium.

## ACKNOWLEDGEMENTS

Special thanks to Katrien Luyten for optimizing the protocols for immunohistochemistry and to Annouschka Laenen for performing the statistical analyses of the behavioral tests. Furthermore, we would like to thank the staff of the animal facility at KU Leuven for taking care of our animals.

## ETHICS STATEMENT

The Belgian and European laws on care and use of laboratory animals were followed. Furthermore, the ARRIVE guidelines were taken into account when preparing this manuscript.

## SUPPLEMENTARY DATA

**Supplementary Table 1:** Detailed information of the protocols used for the different immunohisto-chemical stainings.

| Primary antibody | Dilution | Manufacturer | Antigen retrieval | Time | Visualization staining | Positive control |
| --- | --- | --- | --- | --- | --- | --- |
| **Cytokeratin** | 1/1500 | DAKO (USA) | Pepsin (37°C) | 2 hours at RT | DAB 10 minutes | Murine uterine tissue |
| **ER-alpha** | 1/300 | Santa Cruz (USA) | Citrate | Overnight at 4°C | DAB peroxidase 2 minutes | Murine uterine tissue |
| **Ki67** | 1/3000 | Abcam (United Kingdom) | Citrate | Overnight at 4°C | DAB 10 minutes | Murine skin tissue (paw) |
| **Pgp 9.5** | 1/1000 | DAKO (USA) | Citrate | Overnight at 4°C | DAB 10 minutes | Murine skin tissue (paw) |
| **Podoplanin** | 1/1200 | Abcam (United Kingdom) | Citrate | Overnight at 4°C | NBT 2 minutes | Murine lung tissue |
| **Progesterone** | 1/100 | Santa Cruz (USA) | Citrate | Overnight at 4°C | DAB peroxidase 2 minutes | Murine uterine tissue |
| **Vimentin** | 1/1500 | Abcam (United Kingdom) | Tris-EDTA | 1 hour at 37°C | DAB 10 minutes | Murine kidney tissue |

**Supplementary Table 2:**
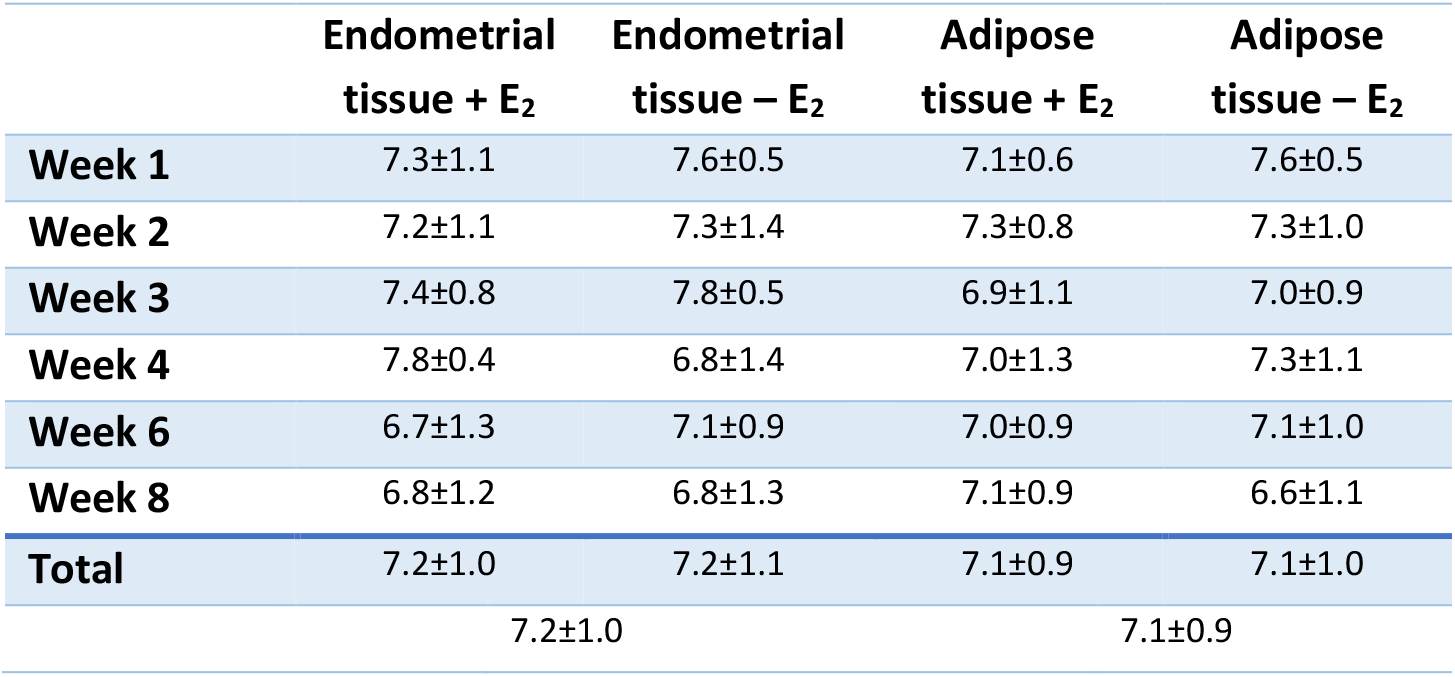
Allred score after PR staining in attached peritoneal implants of recipient mice.

|  | Endometrial tissue + E<sub>2</sub> | Endometrial tissue – E<sub>2</sub> | Adipose tissue + E<sub>2</sub> | Adipose tissue – E<sub>2</sub> |
| --- | --- | --- | --- | --- |
| **Week 1** | 7.3±1.1 | 7.6±0.5 | 7.1±0.6 | 7.6±0.5 |
| **Week 2** | 7.2±1.1 | 7.3±1.4 | 7.3±0.8 | 7.3±1.0 |
| **Week 3** | 7.4±0.8 | 7.8±0.5 | 6.9±1.1 | 7.0±0.9 |
| **Week 4** | 7.8±0.4 | 6.8±1.4 | 7.0±1.3 | 7.3±1.1 |
| **Week 6** | 6.7±1.3 | 7.1±0.9 | 7.0±0.9 | 7.1±1.0 |
| **Week 8** | 6.8±1.2 | 6.8±1.3 | 7.1±0.9 | 6.6±1.1 |
| **Total** | 7.2±1.0 | 7.2±1.1 | 7.1±0.9 | 7.1±1.0 |
|  | 7.2±1.0 |  | 7.1±0.9 |  |

**Supplementary Table 3:**
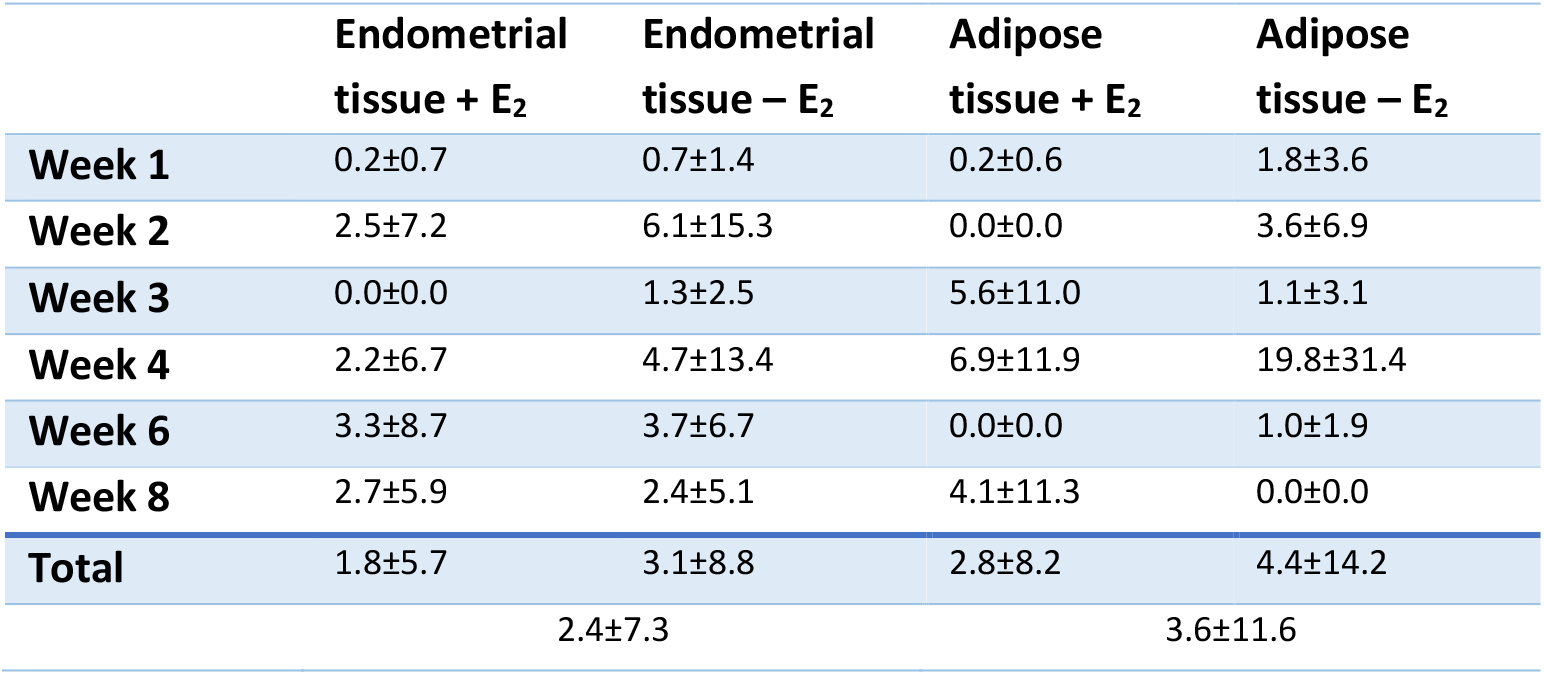
Number of nerve fibers per mm^2^ after PGP9.5 staining in attached peritoneal implants of recipient mice.

|  | Endometrial tissue + E<sub>2</sub> | Endometrial tissue – E<sub>2</sub> | Adipose tissue + E<sub>2</sub> | Adipose tissue – E<sub>2</sub> |
| --- | --- | --- | --- | --- |
| **Week 1** | 0.2±0.7 | 0.7±1.4 | 0.2±0.6 | 1.8±3.6 |
| **Week 2** | 2.5±7.2 | 6.1±15.3 | 0.0±0.0 | 3.6±6.9 |
| **Week 3** | 0.0±0.0 | 1.3±2.5 | 5.6±11.0 | 1.1±3.1 |
| **Week 4** | 2.2±6.7 | 4.7±13.4 | 6.9±11.9 | 19.8±31.4 |
| **Week 6** | 3.3±8.7 | 3.7±6.7 | 0.0±0.0 | 1.0±1.9 |
| **Week 8** | 2.7±5.9 | 2.4±5.1 | 4.1±11.3 | 0.0±0.0 |
| **Total** | 1.8±5.7 | 3.1±8.8 | 2.8±8.2 | 4.4±14.2 |
|  | 2.4±7.3 |  | 3.6±11.6 |  |

**Supplementary Table 4:** Number of lymph vessels per mm^2^ after podoplanin staining in attached peritoneal implants of recipient mice.

|  | Endometrial<br>tissue + E<sub>2</sub> | Endometrial<br>tissue – E<sub>2</sub> | Adipose<br>tissue + E<sub>2</sub> | Adipose<br>tissue – E<sub>2</sub> |
| --- | --- | --- | --- | --- |
| **Week 1** | 0.80±1.68 | 1.57±3.33 | 1.04±1.88 | 0.63±0.95 |
| **Week 2** | 1.70±3.96 | 0.23±0.65 | 0.50±0.93 | 0.40±0.80 |
| **Week 3** | 0.67±1.63 | 1.99±3.32 | 2.50±3.69 | 1.07±1.71 |
| **Week 4** | 8.28±9.29 | 1.17±2.28 | 2.34±5.73 | 3.78±6.75 |
| **Week 6** | 6.05±12.38 | 3.13±7.19 | 2.54±4.68 | 1.16±0.82 |
| **Week 8** | 18.23±41.89 | 0.76±1.41 | 1.82±2.86 | 0.42±1.10 |
| **Total** | 5.98±18.64 | 1.48±3.61 | 1.77±3.42 | 1.29±3.04 |
|  | 3.68±13.41 |  | 1.52±3.22 |  |

**Supplementary Table 5:** Withdrawal threshold of the recipient animals before and after tissue inoculation using the Von Frey hair test.

|  | Endometrial<br>tissue + E<sub>2</sub> |  | Endometrial<br>tissue – E<sub>2</sub> |  | Adipose tissue +<br>E<sub>2</sub> |  | Adipose tissue –<br>E<sub>2</sub> |  |
| --- | --- | --- | --- | --- | --- | --- | --- | --- |
|  | Before | After | Before | After | Before | After | Before | After |
| **Week 1** | 2.4±0.9 | 2.2±0.9 | 2.7±0.7 | 3.0±0.7 | 2.7±0.8 | 2.1±0.7 | 2.7±0.7 | 2.6±0.7 |
| **Week 2** | 2.8±0.8 | 2.4±0.8 | 2.7±0.8 | 2.1±0.6 | 2.3±1.2 | 2.0±0.6 | 2.6±0.8 | 2.0±0.8 |
| **Week 3** | 2.7±0.7 | 2.2±0.8 | 2.4±0.7 | 2.2±0.4 | 2.6±0.7 | 2.0±0.6 | 2.3±0.9 | 2.1±0.9 |
| **Week 4** | 2.1±0.7 | 2.2±0.6 | 2.1±0.6 | 2.1±0.7 | 2.8±0.7 | 2.3±0.8 | 2.6±0.7 | 2.5±1.0 |
| **Week 6** | 2.8±1.0 | 2.4±1.0 | 2.0±0.7 | 2.2±0.6 | 2.2±0.8 | 2.0±0.9 | 2.4±1.0 | 2.2±0.6 |
| **Week 8** | 2.5±0.5 | 2.7±0.9 | 2.6±0.4 | 2.3±0.6 | 2.5±0.7 | 2.3±0.7 | 2.6±0.8 | 2.1±0.7 |

## REFERENCES

1. Giudice, L.C. and L.C. Kao, Endometriosis. Lancet, 2004. 364(9447): p. 1789–99.

2. Sampson, J.A., Peritoneal endometriosis due to the menstrual dissemination of endometrial tissue into the peritoneal cavity. American journal of obstetrics and gynecology, 1927(14): p. 422–469.

3. Giudice, L.C., Clinical practice. Endometriosis. N Engl J Med, 2010. 362(25): p. 2389–98.

4. de Ziegler, D., B. Borghese, and C. Chapron, Endometriosis and infertility: pathophysiology and management. Lancet, 2010. 376(9742): p. 730–8.

5. Schrager, S., J. Falleroni, and J. Edgoose, Evaluation and treatment of endometriosis. Am Fam Physician, 2013. 87(2): p. 107–13.

6. Kennedy, S., A. Bergqvist, C. Chapron, T. D’Hooghe, G. Dunselman, R. Greb, et al., ESHRE guideline for the diagnosis and treatment of endometriosis. Hum Reprod, 2005. 20(10): p. 2698–704.

7. Taylor, H.S., K.G. Osteen, K.L. Bruner-Tran, C.J. Lockwood, G. Krikun, A. Sokalska, et al., Novel therapies targeting endometriosis. Reprod Sci, 2011. 18(9): p. 814–23.

8. Stratton, P. and K.J. Berkley, Chronic pelvic pain and endometriosis: translational evidence of the relationship and implications. Hum Reprod Update, 2011. 17(3): p. 327–46.

9. Howard, F.M., Endometriosis and mechanisms of pelvic pain. J Minim Invasive Gynecol, 2009. 16(5): p. 540–50.

10. Kobayashi, H., Y. Yamada, S. Morioka, E. Niiro, A. Shigemitsu, and F. Ito, Mechanism of pain generation for endometriosis-associated pelvic pain. Arch Gynecol Obstet, 2014. 289(1): p. 13–21.

11. Yan, D., X. Liu, and S.W. Guo, Nerve fibers and endometriotic lesions: partners in crime in inflicting pains in women with endometriosis. Eur J Obstet Gynecol Reprod Biol, 2017. 209: p. 14–24.

12. Tu, F.F., K.M. Hellman, and M.M. Backonja, Gynecologic management of neuropathic pain. Am J Obstet Gynecol, 2011. 205(5): p. 435–43.

13. Morotti, M., K. Vincent, and C.M. Becker, Mechanisms of pain in endometriosis. Eur J Obstet Gynecol Reprod Biol, 2017. 209: p. 8–13.

14. Liu, J., X. Liu, K. Duan, Y. Zhang, and S.W. Guo, The expression and functionality of transient receptor potential vanilloid 1 in ovarian endometriomas. Reprod Sci, 2012. 19(10): p. 1110–24.

15. Greaves, E., K. Grieve, A.W. Horne, and P.T. Saunders, Elevated peritoneal expression and estrogen regulation of nociceptive ion channels in endometriosis. J Clin Endocrinol Metab, 2014. 99(9): p. E1738–43.

16. Greaves, E., A.W. Horne, H. Jerina, M. Mikolajczak, L. Hilferty, R. Mitchell, et al., EP2 receptor antagonism reduces peripheral and central hyperalgesia in a preclinical mouse model of endometriosis. Sci Rep, 2017. 7: p. 44169.

17. D’Hooghe, T.M., C.M. Kyama, D. Chai, A. Fassbender, A. Vodolazkaia, A. Bokor, et al., Nonhuman primate models for translational research in endometriosis. Reprod Sci, 2009. 16(2): p. 152–61.

18. Rogers, P.A., T.M. D’Hooghe, A. Fazleabas, C.E. Gargett, L.C. Giudice, G.W. Montgomery, et al., Priorities for endometriosis research: recommendations from an international consensus workshop. Reprod Sci, 2009. 16(4): p. 335–46.

19. Grummer, R., Animal models in endometriosis research. Hum Reprod Update, 2006. 12(5): p. 641–9.

20. King, C.M., C. Barbara, A. Prentice, J.D. Brenton, and D.S. Charnock-Jones, Models of endometriosis and their utility in studying progression to ovarian clear cell carcinoma. J Pathol, 2016. 238(2): p. 185–96.

21. Story, L. and S. Kennedy, Animal studies in endometriosis: a review. ILAR J, 2004. 45(2): p. 132–8.

22. Brasted, M., C.A. White, T.G. Kennedy, and L.A. Salamonsen, Mimicking the events of menstruation in the murine uterus. Biol Reprod, 2003. 69(4): p. 1273–80.

23. Greaves, E., F.L. Cousins, A. Murray, A. Esnal-Zufiaurre, A. Fassbender, A.W. Horne, et al., A Novel Mouse Model of Endometriosis Mimics Human Phenotype and Reveals Insights into the Inflammatory Contribution of Shed Endometrium. Am J Pathol, 2014. 184(7): p. 1930–9.

24. Peterse, D., M.M. Binda, D.F. O, A. Vanhie, A. Fassbender, J. Vriens, et al., Of Mice and Women: A Laparoscopic Mouse Model for Endometriosis. J Minim Invasive Gynecol, 2017.

25. Greaves, E., H.O.D. Critchley, A.W. Horne, and P.T.K. Saunders, Relevant human tissue resources and laboratory models for use in endometriosis research. Acta Obstet Gynecol Scand, 2017. 96(6): p. 644–658.

26. Pullen, N., C.L. Birch, G.J. Douglas, Q. Hussain, I. Pruimboom-Brees, and R.J. Walley, The translational challenge in the development of new and effective therapies for endometriosis: a review of confidence from published preclinical efficacy studies. Hum Reprod Update, 2011. 17(6): p. 791–802.

27. Dodds, K.N., E.A.H. Beckett, S.F. Evans, and M.R. Hutchinson, Lesion development is modulated by the natural estrous cycle and mouse strain in a minimally-invasive model of endometriosis. Biol Reprod, 2017.

28. Galvankar, M., N. Singh, and D. Modi, Estrogen is essential but not sufficient to induce endometriosis. J Biosci, 2017. 42(2): p. 251–263.

29. Burns, K.A., K.F. Rodriguez, S.C. Hewitt, K.S. Janardhan, S.L. Young, and K.S. Korach, Role of estrogen receptor signaling required for endometriosis-like lesion establishment in a mouse model. Endocrinology, 2012. 153(8): p. 3960–71.

30. Grummer, R., F. Schwarzer, K. Bainczyk, H. Hess-Stumpp, P.A. Regidor, A.E. Schindler, et al., Peritoneal endometriosis: validation of an in-vivo model. Hum Reprod, 2001. 16(8): p. 1736–43.

31. Bergqvist, A., S. Jeppsson, S. Kullander, and O. Ljungberg, Human uterine endometrium and endometriotic tissue transplanted into nude mice. Morphologic effects of various steroid hormones. The American journal of pathology, 1985. 121(2): p. 337–41.

32. Bergqvist, A., S. Jeppsson, S. Kullander, and O. Ljungberg, Human endometrium transplanted into nude mice. Histologic effects of various steroid hormones. The American journal of pathology, 1985. 119(2): p. 336–44.

33. Colette, S., S. Defrere, J.C. Lousse, A. Van Langendonckt, E. Loumaye, and J. Donnez, Evaluation of estrogen treatment in an immunodeficient mouse endometriosis model. Gynecol Obstet Invest, 2009. 68(4): p. 262–8.

34. Fortin, M., M. Lepine, Y. Merlen, I. Thibeault, C. Rancourt, D. Gosselin, et al., Quantitative assessment of human endometriotic tissue maintenance and regression in a noninvasive mouse model of endometriosis. Mol Ther, 2004. 9(4): p. 540–7.

35. Kulak, J., Jr., C. Fischer, B. Komm, and H.S. Taylor, Treatment with bazedoxifene, a selective estrogen receptor modulator, causes regression of endometriosis in a mouse model. Endocrinology, 2011. 152(8): p. 3226–32.

36. Rudzitis-Auth, J., A. Nenicu, R.M. Nickels, M.D. Menger, and M.W. Laschke, Estrogen Stimulates Homing of Endothelial Progenitor Cells to Endometriotic Lesions. Am J Pathol, 2016. 186(8): p. 2129–2142.

37. Burns, K.A., A.M. Pearson, J.L. Slack, E.D. Por, A.N. Scribner, N.A. Eti, et al., Endometriosis in the Mouse: Challenges and Progress Toward a ‘Best Fit’ Murine Model. Front Physiol, 2021. 12: p. 806574.

38. Dorning, A., P. Dhami, K. Panir, C. Hogg, E. Park, G.D. Ferguson, et al., Bioluminescent imaging in induced mouse models of endometriosis reveals differences in four model variations. Dis Model Mech, 2021. 14(8).

39. Greaves, E., M. Rosser, and P.T.K. Saunders, Endometriosis-Associated Pain - Do Preclinical Rodent Models Provide a Good Platform for Translation? Adv Anat Embryol Cell Biol, 2020. 232: p. 25–55.

40. Nunez-Badinez, P., B. De Leo, A. Laux-Biehlmann, A. Hoffmann, T.M. Zollner, P.T.K. Saunders, et al., Preclinical models of endometriosis and interstitial cystitis/bladder pain syndrome: an Innovative Medicines Initiative-PainCare initiative to improve their value for translational research in pelvic pain. Pain, 2021. 162(9): p. 2349–2365.

41. Greaves, E., J. Temp, A. Esnal-Zufiurre, S. Mechsner, A.W. Horne, and P.T. Saunders, Estradiol is a critical mediator of macrophage-nerve cross talk in peritoneal endometriosis. Am J Pathol, 2015. 185(8): p. 2286–97.

42. Silveira, C.G., D. Finas, P. Hunold, F. Koster, K. Stroschein, G.O. Canny, et al., L1 cell adhesion molecule as a potential therapeutic target in murine models of endometriosis using a monoclonal antibody approach. PLoS One, 2013. 8(12): p. e82512.

43. Greaves, E., F. Collins, A. Esnal-Zufiaurre, S. Giakoumelou, A.W. Horne, and P.T. Saunders, Estrogen receptor (ER) agonists differentially regulate neuroangiogenesis in peritoneal endometriosis via the repellent factor SLIT3. Endocrinology, 2014. 155(10): p. 4015–26.

44. Guo, S.W., Y. Zheng, Y. Lu, X. Liu, and J.G. Geng, Slit2 overexpression results in increased microvessel density and lesion size in mice with induced endometriosis. Reprod Sci, 2013. 20(3): p. 285–98.

45. Cousins, F.L., A. Murray, A. Esnal, D.A. Gibson, H.O. Critchley, and P.T. Saunders, Evidence from a mouse model that epithelial cell migration and mesenchymal-epithelial transition contribute to rapid restoration of uterine tissue integrity during menstruation. PLoS One, 2014. 9(1): p. e86378.

46. Lu, Y., J. Nie, X. Liu, Y. Zheng, and S.W. Guo, Trichostatin A, a histone deacetylase inhibitor, reduces lesion growth and hyperalgesia in experimentally induced endometriosis in mice. Hum Reprod, 2010. 25(4): p. 1014–25.

47. Vernon, M.W. and E.A. Wilson, Studies on the surgical induction of endometriosis in the rat. Fertil Steril, 1985. 44(5): p. 684–94.

48. Cason, A.M., C.L. Samuelsen, and K.J. Berkley, Estrous changes in vaginal nociception in a rat model of endometriosis. Horm Behav, 2003. 44(2): p. 123–31.

49. Peterse, D.P., A. Fassbender, D.F. O, A. Vanhie, P. Saunders, J. Vriens, et al., Laparoscopic Surgery: A New Technique to Induce Endometriosis in a Mouse Model. Reprod Sci, 2016. 23(10): p. 1332–9.

50. Ingberg, E., A. Theodorsson, E. Theodorsson, and J.O. Strom, Methods for long-term 17beta-estradiol administration to mice. Gen Comp Endocrinol, 2012. 175(1): p. 188–93.

51. Van Langendonckt, A., F. Casanas-Roux, J. Eggermont, and J. Donnez, Characterization of iron deposition in endometriotic lesions induced in the nude mouse model. Hum Reprod, 2004. 19(6): p. 1265–71.

52. Van Langendonckt, A., F. Casanas-Roux, and J. Donnez, Iron overload in the peritoneal cavity of women with pelvic endometriosis. Fertil Steril, 2002. 78(4): p. 712–8.

53. Arumugam, K., Endometriosis and infertility: raised iron concentration in the peritoneal fluid and its effect on the acrosome reaction. Hum Reprod, 1994. 9(6): p. 1153–7.

54. Allred, D.C., J.M. Harvey, M. Berardo, and G.M. Clark, Prognostic and predictive factors in breast cancer by immunohistochemical analysis. Mod Pathol, 1998. 11(2): p. 155–68.

55. ter Horst, J.P., E.R. de Kloet, H. Schachinger, and M.S. Oitzl, Relevance of stress and female sex hormones for emotion and cognition. Cell Mol Neurobiol, 2012. 32(5): p. 725–35.

56. Rahn, E.J., T. Iannitti, R.R. Donahue, and B.K. Taylor, Sex differences in a mouse model of multiple sclerosis: neuropathic pain behavior in females but not males and protection from neurological deficits during proestrus. Biol Sex Differ, 2014. 5(1): p. 4.

57. Morgan, M.A. and D.W. Pfaff, Effects of estrogen on activity and fear-related behaviors in mice. Horm Behav, 2001. 40(4): p. 472–82.

58. Sanoja, R. and F. Cervero, Estrogen modulation of ovariectomy-induced hyperalgesia in adult mice. Eur J Pain, 2008. 12(5): p. 573–81.

59. Llaneza, D.C. and C.A. Frye, Progestogens and estrogen influence impulsive burying and avoidant freezing behavior of naturally cycling and ovariectomized rats. Pharmacol Biochem Behav, 2009. 93(3): p. 337–42.

60. Schneider, T. and P. Popik, Attenuation of estrous cycle-dependent marble burying in female rats by acute treatment with progesterone and antidepressants. Psychoneuroendocrinology, 2007. 32(6): p. 651–9.

61. Burma, N.E., H. Leduc-Pessah, C.Y. Fan, and T. Trang, Animal models of chronic pain: Advances and challenges for clinical translation. J Neurosci Res, 2017. 95(6): p. 1242–1256.

62. Boyce-Rustay, J.M., P. Honore, and M.F. Jarvis, Animal models of acute and chronic inflammatory and nociceptive pain. Methods Mol Biol, 2010. 617: p. 41–55.

63. Choi, Y., Y.W. Yoon, H.S. Na, S.H. Kim, and J.M. Chung, Behavioral signs of ongoing pain and cold allodynia in a rat model of neuropathic pain. Pain, 1994. 59(3): p. 369–76.

64. Hargreaves, K., R. Dubner, F. Brown, C. Flores, and J. Joris, A new and sensitive method for measuring thermal nociception in cutaneous hyperalgesia. Pain, 1988. 32(1): p. 77–88.

65. Dixon, W.J., Efficient analysis of experimental observations. Annu Rev Pharmacol Toxicol, 1980. 20: p. 441–62.

66. Bonin, R.P., C. Bories, and Y. De Koninck, A simplified up-down method (SUDO) for measuring mechanical nociception in rodents using von Frey filaments. Mol Pain, 2014. 10: p. 26.

67. Njung’e, K. and S.L. Handley, Evaluation of marble-burying behavior as a model of anxiety. Pharmacol Biochem Behav, 1991. 38(1): p. 63–7.

68. Bergqvist, A., S. Jeppsson, S. Kullander, and O. Ljungberg, Human uterine endometrium and endometriotic tissue transplanted into nude mice. Morphologic effects of various steroid hormones. Am J Pathol, 1985. 121(2): p. 337–41.

69. Korbel, C., M.D. Menger, and M.W. Laschke, Size and spatial orientation of uterine tissue transplants on the peritoneum crucially determine the growth and cyst formation of endometriosis-like lesions in mice. Hum Reprod, 2010. 25(10): p. 2551–8.

70. Zamah, N.M., M.G. Dodson, L.C. Stephens, V.C. Buttram, Jr., P.K. Besch, and R.H. Kaufman, Transplantation of normal and ectopic human endometrial tissue into athymic nude mice. Am J Obstet Gynecol, 1984. 149(6): p. 591–7.

71. Fainaru, O., A. Adini, O. Benny, I. Adini, S. Short, L. Bazinet, et al., Dendritic cells support angiogenesis and promote lesion growth in a murine model of endometriosis. FASEB J, 2008. 22(2): p. 522–9.

72. Corona, R., J. Verguts, M.M. Binda, C.R. Molinas, R. Schonman, and P.R. Koninckx, The impact of the learning curve on adhesion formation in a laparoscopic mouse model. Fertil Steril, 2011. 96(1): p. 193–7.

73. Hsiao, K.Y., N. Chang, S.C. Lin, Y.H. Li, and M.H. Wu, Inhibition of dual specificity phosphatase-2 by hypoxia promotes interleukin-8-mediated angiogenesis in endometriosis. Hum Reprod, 2014. 29(12): p. 2747–55.

74. Nothnick, W.B., A. Graham, J. Holbert, and M.J. Weiss, miR-451 deficiency is associated with altered endometrial fibrinogen alpha chain expression and reduced endometriotic implant establishment in an experimental mouse model. PLoS One, 2014. 9(6): p. e100336.

75. Armstrong, G.M., J.A. Maybin, A.A. Murray, M. Nicol, C. Walker, P.T.K. Saunders, et al., Endometrial apoptosis and neutrophil infiltration during menstruation exhibits spatial and temporal dynamics that are recapitulated in a mouse model. Sci Rep, 2017. 7(1): p. 17416.

76. Das, S.K., Cell cycle regulatory control for uterine stromal cell decidualization in implantation. Reproduction, 2009. 137(6): p. 889–99.

77. Patterson, A.L., L. Zhang, N.A. Arango, J. Teixeira, and J.K. Pru, Mesenchymal-to-epithelial transition contributes to endometrial regeneration following natural and artificial decidualization. Stem Cells Dev, 2013. 22(6): p. 964–74.

78. Renjini, A.P., S. Titus, P. Narayan, M. Murali, R.K. Jha, and M. Laloraya, STAT3 and MCL-1 associate to cause a mesenchymal epithelial transition. J Cell Sci, 2014. 127(Pt 8): p. 1738–50.

79. Pelch, K.E., K.L. Sharpe-Timms, and S.C. Nagel, Mouse model of surgically-induced endometriosis by auto-transplantation of uterine tissue. J Vis Exp, 2012(59): p. e3396.

80. Maddern, J., L. Grundy, A. Harrington, G. Schober, J. Castro, and S.M. Brierley, A syngeneic inoculation mouse model of endometriosis that develops multiple comorbid visceral and cutaneous pain like behaviours. Pain, 2022. 163(8): p. 1622–1635.

81. Fattori, V., N.S. Franklin, R. Gonzalez-Cano, D. Peterse, A. Ghalali, E. Madrian, et al., Nonsurgical mouse model of endometriosis-associated pain that responds to clinically active drugs. Pain, 2020. 161(6): p. 1321–1331.

82. Kusakabe, K.T., H. Abe, T. Kondo, K. Kato, T. Okada, and Y. Otsuki, DNA microarray analysis in a mouse model for endometriosis and validation of candidate factors with human adenomyosis. J Reprod Immunol, 2010. 85(2): p. 149–60.

83. Li, Y., M.K. Adur, A. Kannan, J. Davila, Y. Zhao, R.A. Nowak, et al., Progesterone Alleviates Endometriosis via Inhibition of Uterine Cell Proliferation, Inflammation and Angiogenesis in an Immunocompetent Mouse Model. PLoS One, 2016. 11(10): p. e0165347.

84. Bilotas, M., G. Meresman, I. Stella, C. Sueldo, and R.I. Baranao, Effect of aromatase inhibitors on ectopic endometrial growth and peritoneal environment in a mouse model of endometriosis. Fertil Steril, 2010. 93(8): p. 2513–8.

85. Castro, J., J. Maddern, L. Grundy, J. Manavis, A.M. Harrington, G. Schober, et al., A mouse model of endometriosis that displays vaginal, colon, cutaneous, and bladder sensory comorbidities. FASEB J, 2021. 35(4): p. e21430.

86. Chaldakov, G.N., M. Fiore, A.B. Tonchev, M.G. Hristova, V. Nikolova, and L. Aloe, Tissue with high intelligence quotient; Adipose-derived stem cells in neural regeneration. Neural Regen Res, 2009. 4(12): p. 1116–1120.

87. Fliers, E., F. Kreier, P.J. Voshol, L.M. Havekes, H.P. Sauerwein, A. Kalsbeek, et al., White adipose tissue: getting nervous. J Neuroendocrinol, 2003. 15(11): p. 1005–10.

88. Zhou, S., T. Yi, R. Liu, C. Bian, X. Qi, X. He, et al., Proteomics identification of annexin A2 as a key mediator in the metastasis and proangiogenesis of endometrial cells in human adenomyosis. Mol Cell Proteomics, 2012. 11(7): p. M112 017988.

89. Arosh, J.A., J. Lee, D. Balasubbramanian, J.A. Stanley, C.R. Long, M.W. Meagher, et al., Molecular and preclinical basis to inhibit PGE2 receptors EP2 and EP4 as a novel nonsteroidal therapy for endometriosis. Proc Natl Acad Sci U S A, 2015. 112(31): p. 9716–21.

90. Guo, S.W., D. Ding, J.G. Geng, L. Wang, and X. Liu, P-selectin as a potential therapeutic target for endometriosis. Fertil Steril, 2015.

91. Jingwei, C., D. Huilan, T. Ruixiao, Y. Hua, and M. Huirong, Effect of Bushenwenyanghuayu decoction on nerve growth factor and bradykinin/bradykinin B1 receptor in a endometriosis dysmenorrhea mouse model. J Tradit Chin Med, 2015. 35(2): p. 184–91.

92. Tejada, M.A., A.I. Santos-Llamas, L. Escriva, J.J. Tarin, A. Cano, M.J. Fernandez-Ramirez, et al., Identification of Altered Evoked and Non-Evoked Responses in a Heterologous Mouse Model of Endometriosis-Associated Pain. Biomedicines, 2022. 10(2).

93. Morgan, M.A., J. Schulkin, and D.W. Pfaff, Estrogens and non-reproductive behaviors related to activity and fear. Neurosci Biobehav Rev, 2004. 28(1): p. 55–63.

94. Akkaya, T. and D. Ozkan, Chronic post-surgical pain. Agri, 2009. 21(1): p. 1–9.

95. Li, C.Q., J.W. Zhang, R.P. Dai, J. Wang, X.G. Luo, and X.F. Zhou, Surgical incision induces anxiety-like behavior and amygdala sensitization: effects of morphine and gabapentin. Pain Res Treat, 2010. 2010: p. 705874.

96. Sanoja, R. and F. Cervero, Estrogen-dependent abdominal hyperalgesia induced by ovariectomy in adult mice: a model of functional abdominal pain. Pain, 2005. 118(1-2): p. 243–53.

97. Molinas, C.R., O. Mynbaev, A. Pauwels, P. Novak, and P.R. Koninckx, Peritoneal mesothelial hypoxia during pneumoperitoneum is a cofactor in adhesion formation in a laparoscopic mouse model. Fertil Steril, 2001. 76(3): p. 560–7.

98. D’Hooghe, T.M., C.S. Bambra, B.M. Raeymaekers, I. De Jonge, J.M. Lauweryns, and P.R. Koninckx, Intrapelvic injection of menstrual endometrium causes endometriosis in baboons (Papio cynocephalus and Papio anubis). Am J Obstet Gynecol, 1995. 173(1): p. 125–34.

99. Horne, A.W., S.F. Ahmad, R. Carter, I. Simitsidellis, E. Greaves, C. Hogg, et al., Repurposing dichloroacetate for the treatment of women with endometriosis. Proc Natl Acad Sci U S A, 2019. 116(51): p. 25389–25391.

